# Histidine 73 methylation coordinates *β*-actin plasticity in response to key environmental factors

**DOI:** 10.1101/2022.12.16.520803

**Authors:** Adrien Schahl, Louis Lagardère, Brandon Walker, Pengyu Ren, Hugo Wioland, Maya Ballet, Antoine Jégou, Matthieu Chavent, Jean-Philip Piquemal

## Abstract

The functional importance of the methylation of histidine 73 (H73) in actin remains unclear. Focusing on cytoplasmic *β*-actin, present in all mammalian cells, we use molecular dynamics simulations with a polarizable force field and adaptive sampling to examine the effects of H73 methylation. Our results show that methylation enhances nucleotide binding cleft opening, alters allosteric pathways connecting subdomains 2 and 4 (SD2 and SD4) in G-actin, and affects backdoor openings and inorganic phosphate release in F-actin, as validated by biochemical assays. These effects depend on the nucleotide and ions interacting with the actin. Together, our findings reveal how H73 methylation regulates *β*-actin plasticity and integrates environmental cues.

## 1 Introduction

The actin cytoskeleton is involved in numerous cellular functions, such as cell shape maintenance, proliferation, and migration [1]. Over many decades, studies using *α*-actin, the main actin isoform present in skeletal muscle tissue [2, 3] as a model of choice, revealed that actin undergoes large conformational changes as it executes its function [4]. Thus, it is well established that actin monomers assemble to nucleate and elongate filaments [5], which form the basis of the cytoskeleton. Structurally, addition of actin monomers at filament ends induces a transition from the globular actin monomer (G-actin) to a flattened F-actin structure [6]. In the filament, the actin structural plasticity is also important for its ATPase activity. Actin molecules achieve this plasticity through movement of its four distinct subdomains (SD1-4) that change relative orientations in response to nucleotide binding, ATP hydrolysis, and inorganic phosphate (Pi) release [7]. These events, in turn, modulate the mechanical properties of the filament and interactions with many actin binding proteins [8].

In addition to being modulated by ATP and ADP binding [3, 4, 9], and ATP hydrolysis, actin plasticity also depends on other environmental factors, such as the surrounding ions [10]. Furthermore, recent work has highlighted the importance of the methylation of histidine 73 (H73) in affecting filament formation and the rate of ATP hydrolysis in the monomeric form [11]. Although first reported almost six decades ago [12], H73 methylation has remained a biochemical oddity and continues to be somewhat of a functional mystery. Recent identification of SETD3 as the methyltransferase responsible for catalyzing this rare and rarely studied post-translational modification [11, 13] has reignited interest in the role of H73 methylation in actin biology. Previous structural studies showed that H73 is located within a region of actin known as sensor loop (Fig 1-A) [14]. Given this location, and the specific interactions methylated H73 makes with the nucleotide binding cleft located between subdomains 2 and 4 (SD2 and SD4; Fig 1-A), this post-translational modification (PTM) was proposed to be a key regulator that couples ATP hydrolysis and Pi release with filament formation [15]. However, many details of mechanism(s) that are governed by the methylation of H73 in actin remain unknown. Additionally, the precise role of H73 methylation in actin dynamics and plasticity remains incompletely understood.

**Fig. 1.**
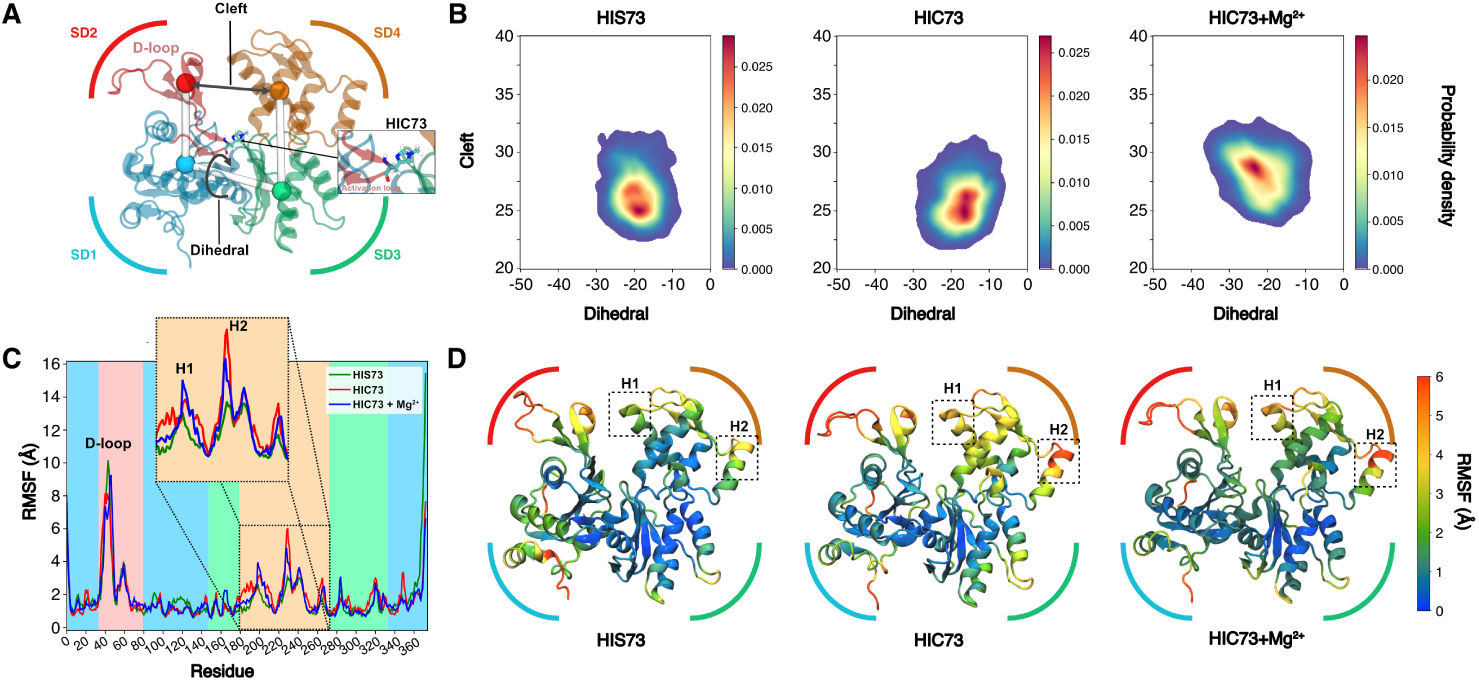
ATP-bound *β*-actin plasticity. **A-** Actin is divided in four subdomains: SD1 (blue), SD2 (red), SD3 (green), and SD4 (tan). The nucleotide-binding cleft (Cleft) represents the distance between SD2 and SD4 domains while the dihedral angle is formed by the four subdomains in the order SD2-SD1-SD3-SD4. The actin is represented with SD2 in upper left and SD4 in upper right (which corresponds to the back of the classic view of actin) to emphasize the positioning of Histidine 73. **B-** Distribution of cleft-dihedrals for actin without methylation (HIS73), or with methylation (HIC73) in presence of KCl and MgCl_2_ (HIC73+Mg^2+^). **C-** RMSF of each system. Zoom on the SD4 subdomain, with a focus on helix H1 (residues 200-206) and helix H2 (residues 228-232). Background colors correspond to subdomains presented in (A). **D-** Representation of each system colored according to their RMSF. Statistical significance of differences have been assessed using a two-sided Kolmogorov-Smirnov test (Table S3)

Here, we address some of these mechanistic questions using *β*-actin as a model system. We chose to focus on *β*-actin because it remains far less studied actin isoform, despite functional relevance. Firstly, unlike *α*-actin, which is only expressed in skeletal muscles, *β*-actin is ubiquitously expressed in all cells. Furthermore, *β*-actin plays a key role in early embryonic development, cell migration and growth [16]. More recently, *β*-actin has been shown to participate in chromatin remodeling [17], and there is accumulating evidence that *β*-actin is important for epithelial-to-mesenchymal transition [18, 19] and of interest in cancer research [20]. However, *β*-actin remains less studied than the *α* isoform, especially in terms of its biochemical mechanism and dynamics. This is primarily due to the fact that obtaining high-quality sample for *in vitro* studies has been a major hurdle for the field, until very recently [21, 22].

Over the last two decades, MD simulations have become an increasingly popular strategy for studying biological systems at different scales [23–27]. Notably, this methodology has been extensively employed to delineate various properties of actin, such as structural differences between nucleotide states [28–30], the localization of water molecules in the actin binding site and their impact on protein plasticity and enzymatic properties [31, 32], interactions with small molecules [33, 34] and the dynamic behavior of filaments in diverse environments [8, 27, 35–41]. However, these computational studies almost exclusively focused on *α*-actin, while the *β*-actin isoform has remained understudied, limiting our understanding of its dynamics and functional implications.

Even if MD simulations have been instrumental in the understanding of how ion and water molecules interact with proteins [42–45] the precise parametrization of these components is still challenging and depends on the system studied [46]. Recent methodological developments on polarizable force-fields have drastically increased the accuracy of interactions between these molecules and proteins [47–57]. In addition, the use of enhanced sampling methods allows exploring diverse unknown states hard to reach through the use of classical molecular dynamics simulations [58–62].

Here, we employed MD simulation strategies that overcome the limitations of classical approaches by combining adaptive sampling with the AMOEBA polarizable force field to more accurately model interatomic interactions. We applied them to provide insights into *β*-actin biochemistry and how H73 methylation together with key environmental factors (ions and nucleotides), affect its dynamics and plasticity both in its core and extremities. This work sheds light onto the plasticity of the *β*-actin isoform, where subtle structural changes can impact actin function, from the monomer to the filament.

## 2 Results

### Histidine 73 methylation modulates the plasticity of ATP bound G-actin

We performed 1.52 µs long adaptive sampling polarizable molecular dynamics simulations for both *α*- and *β*-actin isoforms containing ATP bound nucleotide and magnesium ion in the active site, in the presence of 150 mM of KCl in solution. To assess the effect of H73 methylation on the G-actin dynamics, we simulated both non-methylated histidine (HIS73) and methylated histidine (HIC73) and measured the dihedral angle formed by the four subdomains, as well as the distance between SD2 and SD4 which defines the size of the nucleotide binding cleft [29] (hereafter referred as “cleft”) (Fig. 1A). For HIS73 *β*-actin, the protein fluctuated around a dihedral angle of c.a. −20° and a cleft size of 25 Å (Fig. 1B). The region of high density extends up to 27 Å indicating the capacity of the protein to widen the cleft during the course of the simulation (Fig. S1 and Movie S1). Remarkably, despite differing from HIS73 slightly (presence of an additional -CH3 group), HIC73 induced notable changes in the cleft dynamics, causing an expansion from 25 Å to 27 Å (Fig. S1 and Movie S2), alongside a slight reduction in the dihedral angle to −18°. To assess, which parts of the actin were most affected, we performed Root Mean Square Fluctuations analysis (RMSF) (see Methods) (Fig. 1C). We observed that the D-loop and SD4 subdomain showed distinctive differences in their RMSF profile. More precisely, the dynamics of helices encompassing residues 200-206 (H1 helix) and 228-232 (H2 helix), positioned at opposite sides of SD4, exhibited pronounced sensitivity to histidine methylation (Fig. 1D). Thus, taken together, this suggests that histidine 73 methylation impacts not only the local dynamics around the histidine residue but also regions distal from this site. Interestingly, although the sequence identity between *α*- and *β*-actin is very high (ca 94%, Fig. S2), histidine methylation seemed to affect differently the amplitude of the dihedral-cleft fluctuations for *α* or *β* isoform (Fig. S3). The most affected areas in the *α*-actin were comparable to the ones seen in *β*-actin (Fig. S4). Furthermore, both the dihedral angle and the size of the cleft exhibited a wider range of values in comparison to the *β*-actin isoform. This implies that *β*-actin undergoes more subtle changes compared to the *α*-actin isoform. These results highlight that slight modifications in the amino acid sequence between actin isoforms can be translated into detectable structural variations which may have impact on actin functions. For subsequent sections, we focus on the *β*-actin isoform, as limited information exists on the dynamics of this protein, despite of its importance (as introduced above).

### Histidine 73 methylation opens the SD2-SD4 cleft and affects the enzymatic site

Next, we focused our analysis on dissecting specific structural perturbations in ATP-bound monomeric *β*-actin caused by histidine 73 methylation. We observed that in the case of unmethylated HIS73, residues of SD4 H1 helix interacted with residues from the SD2 domain, effectively bridging the two subdomains (Fig. 2A, left panel). Notably, GLU207 of SD4 formed stable interactions with residue ARG62 of SD2. Given the charged nature of these residues, these interactions are likely mediated via electrostatics. Additionally, based on the bond lengths and angles we measured, hydrogen bonding may also play a role. Regardless of the exact nature of this interaction, when we examined the same interactions in monomeric *β*-actin where HIS73 was methylated (HIC73), we noted that the likelihood of interaction between these amino acids was reduced (Fig. 2A, central panel), facilitating the opening of the cleft. This cleft opening was correlated with an increase of the active site volume (Fig. 2B-C) from 685 Å3 to 760 Å3. As this volume change may affect molecules inside the active site, we scrutinized the dynamic properties of the magnesium ion and water molecules around it. We observed two magnesium ion populations (Fig. 2D): a Mg^2+^ bound to the *γ*-phosphate, with a distance 2.2 Å between these two atoms, and an unbound Mg^2+^ at a distance of 4 Å from the *γ*-phosphate. For HIC73, the population of the Mg^2+^ in the bound state decreased. Surprisingly, we did not observed significant changes in the number of water molecules in the active site (Fig. 2E). Consequently, our results showed how the HIS73 methylation can affect subdomain interactions and active site for ATP-bound *β*-actin.

**Fig. 2.**
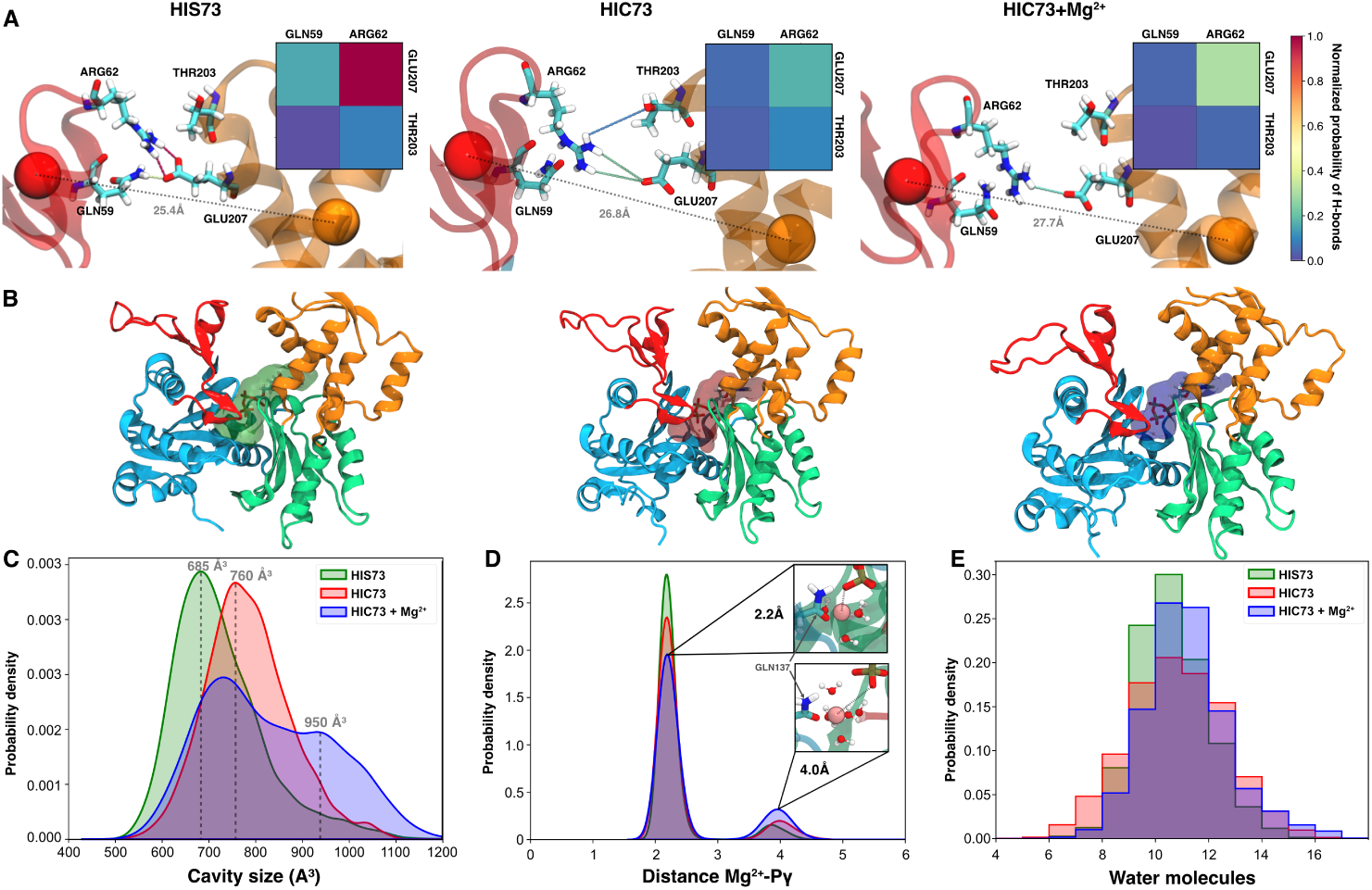
Role of methylation and magnesium ions on ATP-bound *β*-actin intrinsic flexibility. **A-** Representation of main residues interactions at the interface of SD2 and SD4. Interactions are colored in function of the normalized probability of hydrogen bonds between these residues during the simulations. **B-** Volume of the actin active for the most probable actin configurations presented in Figure 1-B. **C-** Distribution of the active site volume for each system presented in (B). **D-** Distribution of the distance between magnesium ion and *γ*-phosphate. **E-** Histograms of the number of water molecules around the magnesium (*≤* 5 Å) in the active site. Statistical significance of differences have been assessed using a two-sided Kolmogorov-Smirnov test (Table S3)

### Histidine 73 methylation exerts allosteric effects to bridge SD2 and SD4 subdomains

Histidine 73 is situated at the core of the actin at the interface of all four subdomains (Fig. 1A). However, the mechanism through which the methylation of the residue influences distant regions, such as helices H1 and H2 on SD4, remains elusive. To elucidate this, we conducted correlation-based dynamical network analysis [63, 64]. This analysis allows calculations of dynamic networks where nodes represent the alpha-carbons of each residue and the edges represent correlations of motions between nodes, and has been used to identify possible, functionally relevant allosteric pathways [64]. Using this approach we identified several possible allosteric pathways between GLU207 and ARG62, the two residues engaged in the hydrogen bond that bridges SD2 and SD4 subdomains (Fig. 2A, left panels). In unmethylated monomeric *β*-actin (HIS73), two most represented allosteric paths involved direct interactions between GLU207 and ARG62 (Fig. 3A, left and central panels; Table S2), while the third path involved additional residues located on both SD2 and SD4 (Fig. 3A, right panel). Interestingly, the sensor loop was directly involved in this path through PRO70 and GLU72. Upon histidine methylation (HIC73 case), the direct interaction between GLU207 and ARG62 was lost, and replaced by interactions among a larger number of residues on SD2 and SD4 subdomains (Fig. 3B and Table S2). Notably, the sensor loop was involved in the two most representative paths, while the third one displayed an SD2-SD4 bridge mediated through interactions between TYR69 and ARG183 (Fig. 3B, right panel). Thus, change from HIS to HIC introduces measurable changes in allosteric network. We also observed that the ARG183 residue was involved in the most represented allosteric path in both HIS73 *β*-actin and HIC73 *β*-actin. To examine further whether a change in a single residue can introduce changes in allosteric networks, we performed simulations of the ARG183GLY mutant for HIC73 *β*-actin. We noted that in the ARG183GLY *β*-actin mutant the paths connecting SD2 and SD4 were redirected around the active site (Fig. S8B-D), which is distinct from what we saw in HIC73 *β*-actin, where they pass through the sensor loop. This rerouting was accompanied by a change of the cleft-dihedral profile (Fig. S8A). Taken together, these findings suggest that histidine 73 methylation promotes the opening of the cleft between SD2 and SD4 subdomains, which is otherwise held in a more closed conformation by the GLU207-ARG62 interaction. Histidine methylation, which changes the local polar environment, also alters allosteric paths that mediated interactions between SD2 and SD4, by rerouting allosteric communications away from direct hydrogen bonding between GLU207 and ARG62 and towards a more significant involvement of the sensor loop.

**Fig. 3.**
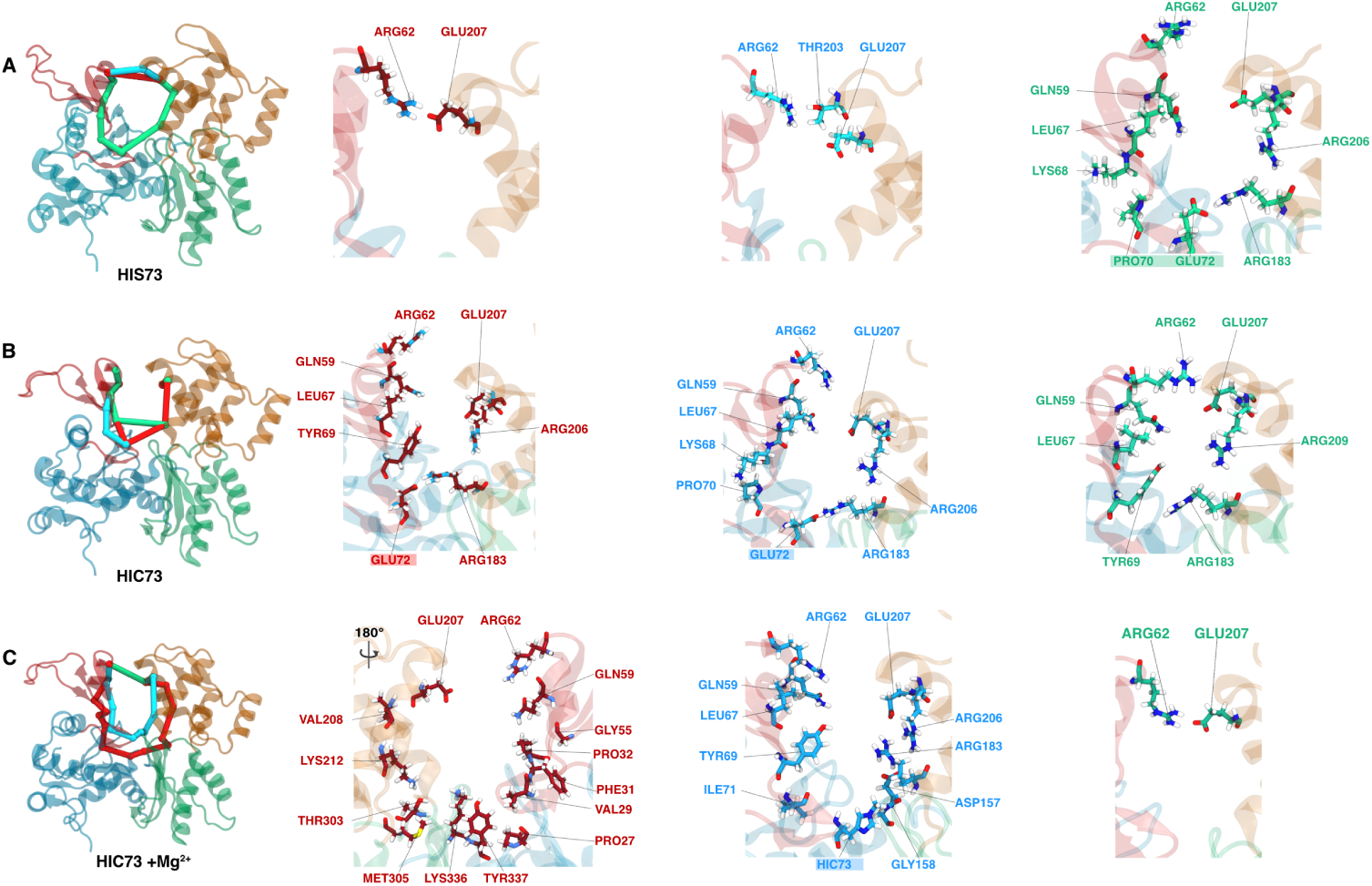
Main allosteric paths in ATP-bound *β*-actin systems. Representation of the three most represented between R62 and E207 for **A-** HIS73, **B-** HIC73 and **C-** HIC73+Mg^2+^ systems. Most represented path in red, second most represented in blue, and third most represented in green. On right, licorice representations of these three main paths using the same color scheme. Residues on the sensor loop are highlighted.

### Presence of magnesium ions modulates the impact of histidine 73 methylation

Magnesium ion concentration plays an important role in actin polymerization [10]. To examine whether histidine methylation is affected by magnesium ions we have performed simulations on the monomeric HIC73 *β*-actin using 75 mM of MgCl2 (to directly compare with 150 mM KCl). At this high Mg^2+^ ion concentration, both cleft and dihedral angles were affected, displaying an increase (up to 30 Å in cleft opening, and up to −30° dihedral angle, respectively) (Fig. 1B, right panel) in comparison to HIC73 system without magnesium ions. The RMSF (Fig. 1C) as well as the volume of the ATP binding site were affected by the addition of magnesium ions. The latter displayed a larger volume (Fig. 2C), although the distribution of water molecules in the active site remained unaffected (Fig. 2E). Additionally, the population of magnesium ion not bound to the *γ*-phosphate slightly increased (Fig. 2D). We also used a larger simulation box of c.a. 100,000 atoms (see Table S1) to simulate a system with a molar concentration of 7.5 mM MgCl2 (corresponding to 5 Mg^2+^ ions), which is similar to the concentration used for *in vitro* actin polymerization assays [10]. At this Mg^2+^ ion concentration, we also observed increases in both cleft and dihedral angles (Fig. S5), albeit less pronounced than the one seen for 75 mM of MgCl2. Attempting a concentration of 75 mM of MgCl2 for this larger system size, we observed even larger modulations of the cleft-dihedral profile (Fig. S5). Thus, monomeric actin flexibility is dependent of the magnesium ion concentration. Magnesium (as well as potassium) ions were interacting on numerous sites spread on different subdomains (ASP56, GLU93, GLU99, GLU100, GLU270, and GLU364; Fig. S6). Lastly, we examined how magnesium ions affect the allosteric paths between SD2 and SD4 in methylated G-actin (Fig. 3C). Interestingly, the most represented path was no longer passing through the sensor loop, but on the opposite side of the active site. Hence, the addition of magnesium ions modulates the effects of H73 methylation and rebalances allosteric pathways in a way that mirrors what we observed in unmethylated G-actin.

### ADP binding reduces the impact of histidine 73 methylation on actin dynamics

To investigate whether the effects of HIS73 methylation are dependent on the nature of the bound nucleotide, we performed additional simulations with ADP in the binding cavity, providing a basis for comparison with ATP-bound actin. In the presence of ADP, the internal flexibility of the actin monomer, particularly in the SD4 subdomain, was reduced for methylated H73, as revealed by the RMSF profile (Fig. 4A), although the dynamics of the H2 helix (residues 228-232) still remained enhanced. Additionally, when compared to ATP-bound state, ADP-binding led to decrease in the cleft opening in HIC73, and HIC73 with addition of Mg^2+^ (Fig. 4B). This difference can be explained by sustained interactions between residues at the SD2-SD4 interface (Fig. 4C and Fig. S7). We observed that when ADP is bound, the volume of the active site exhibited reduced fluctuations (Fig. 4D). In all our conditions (HIS73, HIC73, HIC73+Mg^2+^), the magnesium ion in the binding site was directly bound to the nucleotide (Fig. 4E), despite variations in the number of water molecules around the ion (Fig. 4F). Finally, the binding of ADP also affected the allosteric paths between SD2 and SD4 by involving a direct interaction between ARG62 and GLU207 (Fig 4. G,H and Fig. S7), or by passing through TYR69-ARG183. Thus, in the conditions observed in our simulations, the binding of ADP reduced the impact of histidine 73 methylation on actin plasticity.

**Fig. 4.**
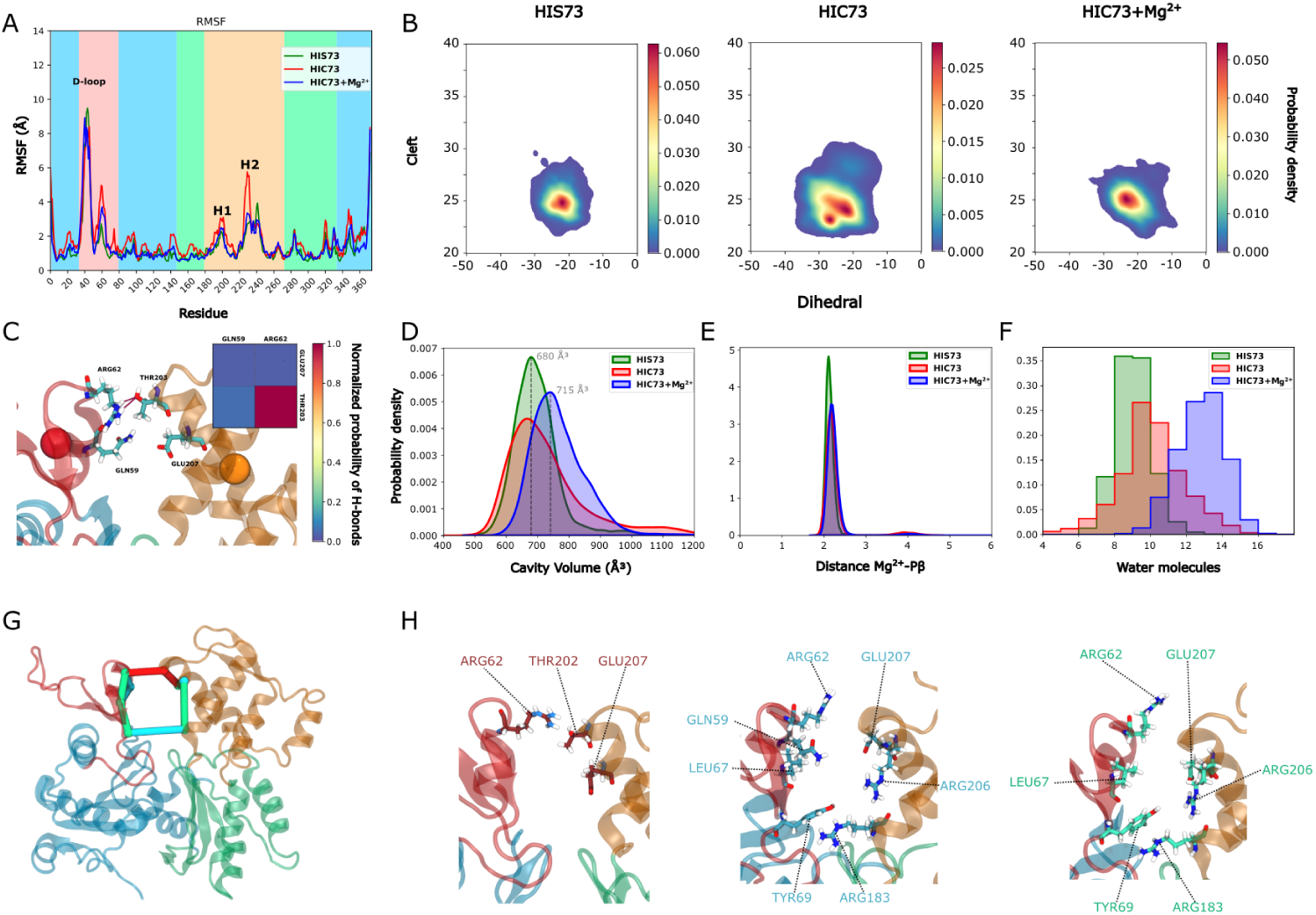
ADP reduces the *β*-actin dynamics. **A-** RMSF of ADP-Actin systems. Background colors correspond to subdomains (see Figure 1). **B-** Cleft-Dihedral maps of ADP bound actin systems. **C-** Main hydrogen bonds between SD2 and SD4 subdomains for HIC73 system. **D-** Volumes of active sites. **E-** Distribution of the distance between magnesium ion and *γ*-phosphate. **F-** Histograms of the number of water molecules around the magnesium (*≤* 5 Å) in the active site. **G-** Representation of three most represented allosteric paths between R62 and E207 in HIC73 system. **H-** Licorice representation of the residues involved in the three main paths presented in G. Pathways for HIS73 and HIC73+Mg^2+^ systems are presented in Figure S4. Statistical significance of differences have been assessed using a two-sided Kolmogorov-Smirnov test (Table S3)

### Impact of the deafness-causing mutation on histidine 73 methylated actin dynamics

The mutation of lysine 118 into asparagine (LYS118ASN) has been shown to cause deafness [65], most likely due to its impact on the rate of actin polymerization and nucleation [66, 67]., To have a better comprehensive view on the influence of this mutation on actin filament nucleation, we conducted simulations with LYS118ASN actin in the presence of both ATP or ADP and HIS73 methylation (HIC73). In the presence of ATP, the cleft-dihedral profile (Fig. S3) revealed a more open state, yet more focused, closely resembling conformations seen in the case of HIC73 with addition of magnesium ions. Interestingly, for bound ATP, the actin flexibility was also reduced (Fig. S4) compared to the wild type HIC73. This suggests that the LYS118ASN mutation not only induces a more open conformation but also imparts a degree of structural rigidity to the actin filament in the presence of ATP. Conversely, in the presence of ADP, the mutation had a very low impact on the cleft-dihedral map (Fig. S3). Interestingly, theses results highlight that even residues that are remote from the nucleotide binding pocket do influence the nucleotide state-dependent flexibility of G-actin, with consequences on nucleation and polymerization kinetics [66, 67].

### Histidine 73 methylation stabilizes F-actin at the barbed end

Transition from G-actin to F-actin that accompanies actin polymerization is essential for cellular function. To understand the impact of H73 methylation on F-actin assembly and dynamics, as a function of nucleotide binding, we analyzed conformational dynamics of actin subunits at the barbed end of the filament. Simulations were performed on the four F-actin subunits, with the two subunits towards the pointed end constrained to mimic a longer filament, and two on the barbed (growing) end free to sample broad conformational space (see details in the Method section). For all the subunits, H73 was either methylated (HIC73) or non-methylated (HIS73), and the active site contained either ATP or ADP. Utilizing 1.52 µs of polarizable MD simulations with adaptive sampling, we explored the dynamics of the last (B) and penultimate (B-1) subunits at the barbed end (Fig. 5A, Fig. S9-S11), which have been reported to experience major deformations [36]. In the methylated system, the last F-actin (B) subunit exhibited more pronounced deformation when ADP was bound, than in the presence of ATP (Fig. S10A), which is in contrast to what we observed in G-actin. Specifically, with ADP bound, the cleft-dihedral profile of HIC73 (see Fig. S11) displayed conformations going towards those observed for HIS73 G-actin monomers (Fig. 4B). On the contrary, with ATP bound, the B subunit was stabilized in a more flatten filament-like form (Fig. S11) as seen in recent cryo-EM structures of the barbed end [8, 68]. The penultimate (B-1) subunit appeared less affected by the nucleotide state, probably due to a higher number of contacts with the other subunits in the filament (Fig. S9 and Fig. S10B). For both B and B-1 subunits, the dynamics of the H1 helix was the most affected in comparison to the monomeric form (Fig. S10). For HIC73, in the B subunit, this helix strongly interacted with the C-terminal residues of the B-1 subunit for bound ADP but not in the case of ATP (Fig. S12). Both ATP- and ADP-bound HIS73 systems were less stabilized in the F-actin conformation than the ATP-bound HIC73 system (Fig. 5B). We also observed more interactions between the C-terminal part of the B-1 subunit and the H1 helix of the B subunit in these systems (Fig. S12). These results showed that H73 methylation also modulated multiple different aspects of F-actin plasticity, further highlighting the biological significance of this PTM.

**Fig. 5.**
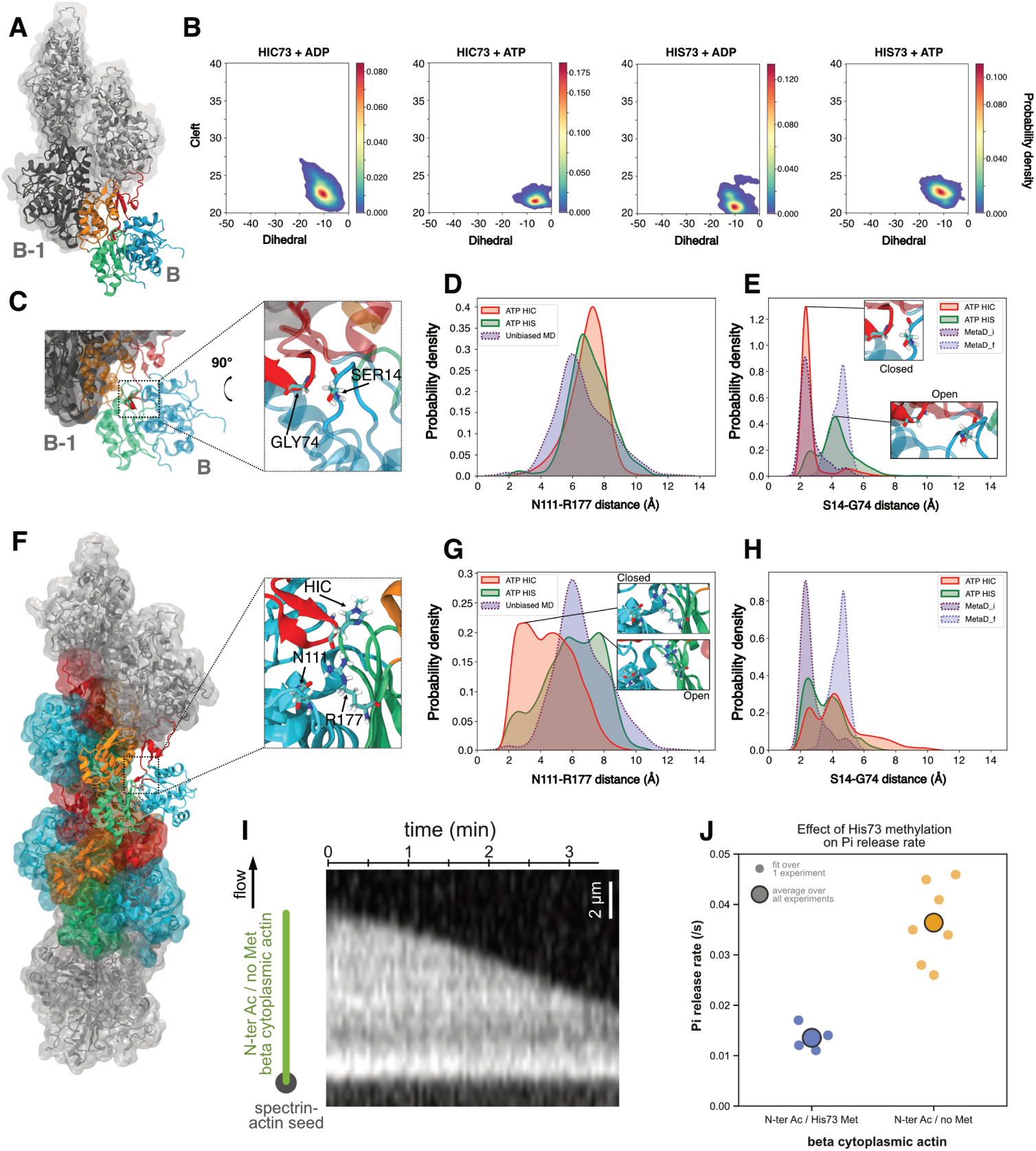
Actin flexibility at the barbed end and inorganic phosphate release. **A-** Representation of ultimate (subunit B, colored) and penultimate (subunit B-1, dark grey) at the filament barbed end. **B-** Distribution of Cleft-Dihedrals angles for ultimate (B) subunit in HIC73 and HIS73 systems containing either ADP or ATP molecules. “Statistical significance of differences have been assessed using a two-sided Kolmogorov-Smirnov test (Table S3)”. **C-** Zoom on the G74-S14 backdoor in the ultimate subunit. **D-Distribution of distances between N111 and R177 residues in HIS73+ATP and HIC73+ATP states of B subunit. The blue curve corresponds to distances extracted from unbiaised MD simulations** [8]**. E-** Distribution of distances between S14 and G74 residues in HIS73+ATP and HIC73+ATP states of B subunit. The metaD i (purple) and metaD f (blue) distributions correspond respectively to distances observed in the initial and final states of metadynamics simulation [8] (see Method). In this simulation the G74-S14 backdoor is closed in the first half of the simulation(metaD i, purple dashed curve), and open in the second half of the simulation (metaD f blue dashed curve). **F-** Representation of the core subunits, with a zoom on the ASN111-ARG177 backdoor. Light grey subunits were harmonically restrained during the simulations. **G-** Averaged distribution of distances between N111 and R177 residues in HIS73+ATP and HIC73+ATP states in the core subunits. **H-** Averaged distribution of distances between G74 and S14 residues in HIS73+ATP and HIC73+ATP states in the core subunits. **I-** (left) In vitro actin filament depolymerization assay performed in microfluidics. Fluorescently-labeled *β*-actin filaments are exposed to buffer only to induce their depolymerization (see Methods). (right) Kymograph of a depolymerizing actin filament. **J-** Pi release rates measured experimentally for HIC73 or HIS73 *β*-actin filaments. Individual data points represent the fitted value of the Pi release rate, for at least 30 filaments per assay. Large data points are the mean values of those repeated experiments. Statistical significance of differences have been assessed using T-test: p-value = 1.1 10^−4^.

### Histidine 73 methylation reduces the opening of molecular backdoors at the barbed end and in the core of the filament

Molecular dynamics simulations were recently used to propose two plausible pathways for the release of inorganic phosphate (Pi) after ATP hydrolysis in the F-actin state [8]. The first pathway involves the breaking of the interaction between ARG177 and ASN111, and was experimentally validated, while the second one requires breaking of the SER14-GLY74 interaction (see Fig. 5C) [69]. This pathway remains to be experimentally demonstrated. Based on our simulations, this second backdoor was mainly closed in the presence of methylated histidine, and open for non-methylated histidine in the ATP-bound state of both B (Fig. 5D,E) and B-1 (Fig. S13) actin subunits. Interestingly, the reanalysis of the data openly deposited by the Hummer lab [8] (see Method section) agreed well in term of opening and closing distances for this second backdoor (Fig. 5D,E S13). Additionally, Oosterheert and coauthors proposed that the ARG177-ASN111 backdoor was open for the last actin subunit [8]. We also observed the opening of this backdoor in the last subunit, independently of the histidine methylation (Fig. S13).

We have also performed 400 ns simulations of the filament actin core using eight F-actin subunits containing ATP. The two subunits at the pointed and barbed extremities were constrained to mimic an infinite filament (see Methods and Fig. 5-F). We analyzed the dynamics of the four central subunits. In average, the ARG177-ASN111 backdoor was mainly open for the non methylated HIS73, while it was mainly closed for the methylated HIC73 (Fig. 5-G). In our simulations, the SER14-GLY74 backdoor display both open and close states for unmethylated and methylated histdine 73 (Fig. 5-G). Analyzing each subunit separately (Fig. S14) highlighted some heterogeneity in terms of Cleft-Dihedral and backdoor opening. This heterogeneity may be due to the constrains applied at the extremities of our model to mimic an infinite filament. Nevertheless, we can notice that the ARG177-ASN111 backdoor was more open for actin subunits in a configuration closed to the F-actin for both methylated and unmethylated histdine 73 (Fig. S14).We also looked at the H161 *ω* angle (Fig. S14), which may adopt two different values, 43.2° in F-form and −85.6° in G-form [8], and has been recently identified as potentially important for Pi release [70]. In our simulations this angle may adopt both values depending on considered subunit. Longer simulations on larger systems may help refining these trends. Taken together, our results suggest a faster Pi release for actin filaments containing unmethylated HIS73 due to backdoors opening in the core of the filament and at the barbed end.

To test the impact of the histidine methylation on the release of Pi within F-actin subunits, we performed *in vitro* actin filament depolymerization assays using microfluidics [71] (Fig 5-I, see Methods). We observed that the Pi release rate was almost 3-fold lower for histidine-methylated *β*-actin compared to the non-methylated one (Fig. 5J and Fig. S15). Thus, our data, together with the recent observations [8], indicate that histidine methylation controls the release of Pi in F-actin, mainly by limiting the opening of the ARG177-ASN11 backdoor in the core of the filament, and potentially of the SER14-GLY74 backdoor at the barbed end.

## 3 Discussion

In this study, we performed adaptive sampling simulations using a polarizable force field on 19 different actin systems (Table S1), reaching a total simulation time of more than 26 µs. Although actin has been extensively studied using classical computational strategies [27–30, 30–32, 36, 45] in this work we used a polarizable force field, which improves accuracy of modeling interatomic interactions. This accuracy, in turn, enables more precise insights into molecular determinants of conformational changes and dynamics that accompany actin assembly and function. Notably, the models of the barded end and core filament, containing respectively from 250k to 525k atoms, represent, to the best of our knowledge, ones of the largest systems being modeled with this type of force field and for such extended timescale. This highlights that MD simulations in combination with polarizable force fields can now be used on large molecular systems to study biologically relevant mechanisms. Due to the size of the actin filament system, we only performed 400 ns of simulations. In the future, longer simulations on larger systems (e.g. filament containing 12 to 16 monomers) will help obtaining more sampling to refine the results presented here. With the continuous growths of computing ability with graphics processing units (GPUs) or exascale supercomputers [72], this type of modeling will become more common.

Although focusing our core interest on *β*-actin, as a more prevalent yet less understood actin isoform, we used *α*-actin to benchmark our approach. Overall, our results for this isoform are in broad agreement with previous atomistic simulations [29, 30], supporting the validity of polarizable force field approach. For example, in our study the cleft distance varied from 23 to 25 Å in the ATP state, which is similar to what was seen by Saunders et al. [29]. However, we note that the dihedral angle values varied among the studies: −18 to −23 degrees for Saunders *et al.* [29], −11.7 degrees in recent work by Jepsen et al. [30], and −12 to −18 degrees in our calculations. Some of this variability may be due to the intrinsic difference between the results obtained using of a polarizable force field and those from non-polarizable atomistic force field simulations, as recently demonstrated for other proteins [52]. Importantly, our simulations revealed that *α*- and *β*-actin exhibited distinct patterns of flexibility (see also Fig. S4 for a direct RMSF comparison), suggesting that, despite the high sequence identity between isoforms, mammals do use different isoforms to build specific networks tailored to specific functions. Amino acid difference from *α*-actin to *β*-actin are spread on three subdomains (see Fig. S2). These structural differences may modulate allosteric paths directly or indirectly. Thus, it would be informative, in a future study, to perform in silico mutations for these specific residues to see how each one individually affects *α*- or *β*-actin dynamics. Our observations on the deafness-causing K118N actin mutant already suggests how actin processes, such as filament nucleation and elongation, are sensitive to actin monomer flexibility. Differences observed in our simulations are consistent with a difference in phosphate release rate between *α*-actin (in between 0.008/s and 0.01/s - see Fig. S15) and *β*-actin (*>* 0.01/s - see Fig. 5-J), the latter releasing its phosphate slightly faster [71] (Fig. S15). Therefore, our study may point at protein dynamics as the key difference between otherwise almost identical actin isoforms expressed by mammals.

In addition to expanding our understanding of *β*-actin, we also addressed the role of a PTM which is highly conserved in mammals, H73 methylation, whose function is yet incompletely understood [11, 13]. Recent studies have shown that unmethylated actin shows an increased rate of nucleotide exchange in the monomeric form, and a slower filament assembly rate [11, 73]. Additionally, H73 methylation was shown to increase actin filament stability [13]. Our results reveals an enhanced internal dynamics of the methylated actin monomer in the ATP state, which may account for these previous observations. Our simulations of the *β*-actin isoform highlighted a wider opening of the cleft between SD2 and SD4 subdomains in the G-actin form due to the disruption of direct interactions between ARG62 and GLU207. This opening may have implications during the early steps required for the nucleation of actin filaments. As ARG62 and GLU207 residues are also involved in inter-subunits interactions in the filament, they thus need to be rearranged accordingly [6]. Based on our results, we speculate that direct interactions between ARG62 and GLU207 may affect interactions between actin subunits, thereby limiting the nucleation process.

We also examined mutual relationships between H73 methylation and environmental factors, such as magnesium ions and nucleotides. In general, we observed that actin plasticity is affected not only by H73 methylation, but by other factors as well. Most notably, ADP binding may lock the actin monomer in an closed state, impeding G-actin structural adaptation at the filament barbed end. It is therefore reasonable to think that interactions between intra-monomer residues may compete with interactions between inter-monomer residues both during the nucleation and elongation of filaments. Our results demonstrate how histidine methylation affects these essential interactions. Finally, histidine methylation may also play a role in stabilizing subunits at the barbed end, as in the absence of methylation, the last subunit tends to relax more quickly towards the G-actin form

Using correlation-based dynamical network analysis we were able to link actin flexibility with distinct allosteric paths between the SD2 and SD4 subdomains of *β*-actin, which are critical for defining the size and openness of the nucleotide binding cleft, and thus ATP hydrolysis. This led to the insight that in the case of ATP-bound methylated G-actin, the main allosteric paths were re-routed through the sensor loop. Conversely, allosteric paths observed in the ADP-bound states almost never involved this loop, minimizing the impact of histidine methylation on SD2-SD4 mobility. Altogether, these results offer initial insights into how allosteric paths drive local molecular rearrangements in the active site towards large conformational changes at the actin filament barbed and minus ends, and how they may fine-tune mechanical properties of actin filaments [9, 74]. This may further inform our understanding of why side-binding proteins, such as ADF/cofilin, are highly sensitive to both the nucleotide state of actin filaments and their mechanical state [75–78].

Our results also highlighted that H73 methylation increases the volume of the active site, without changing the number of water molecules inside the active site. This may indicate that methylation influences the structuring of the water network that participates in ATP hydrolysis [32, 79], rather than the amount of water molecules available. Our results also highlight that histidine methylation tunes Pi release. The Pi release rate within filaments is faster for non-methylated *β*-actin, which may be related to the opening of more than one backdoors, as further revealed by our simulations of core subunits as well as ones located at the filament barbed end. Wang and co-authors recently proposed that the cleavage of interactions between Mg^2+^ and *γ*Pi is the limiting step of Pi release [70]. They also showed that the number of water molecules coordinating the Pi had to increase prior to dissociation with Mg^2+^. Regarding this, the opening of the N111-R177 backdoor might allow the entry of water molecules in the cavity, hence potentially increasing the solvation of the Pi.

Overall, our results highlighted how the post translational modification of histidine 73 can drastically change the dynamic properties of actin in different forms, from the monomer to the filament. Our work reveals how subtle structural changes, such as a single methylation buried within the core of the protein, could have numerous consequences that affect protein plasticity at different scales. Importantly, although understudied, histidine methylation has now been reported in other proteins, such as S100A9, myosin, skeletal muscle myosin light chain kinase (MLCK 2), and ribosomal protein Rpl3 [80], with a recent analysis identifying about 300 histidine methylation sites in the proteome of HeLa cells [81]. Similar to what we observed in *β*-actin, we expect that this PTM plays a major regulatory role in many of these systems. Thus, our study highlights the value of using MD simulations combined with polarizable forcefields as a method for understanding the effects of this PTM in actin and beyond.

## 4 Methods

### Systems preparation for Molecular Dynamics simulations

The initial structures of the mammalian *α*-skeletal actin monomer bound to ATP or ADP were based on the 1NWK [14] and 1J6Z [82] pdb files respectively. For the *β*-actin, we build an homology model by replacing the residues of interest in the structure of the *α*-actin using the ACTB sequence (see Fig. S2). As the N-terminal acetylation or arginylation of the *β*-actin is known to have diverse biological effects [3, 83], we decided to remove the N-terminal amino acid, starting our *β*-actin sequence at ASP2. The N-terminal and C-terminal extremities and the D-loop were generated using modeller [84]. In the original pdb files, the nucleotide is under the form of AMPPNP and in presence of a calcium ion. We replaced the AMPPNP by ATP and the calcium ion by a magnesium ion, to be closer to physiological conditions.

It has been demonstrated that crystallographic water molecules located inside the cavity may impact the behaviour of the protein [31]. Therefore, the water molecules placed at less than 10Å of the magnesium ion were kept.

In addition to the monomer systems, two 4-mer systems and 2 8-mer systems were prepared based on the 6BNO [85] pdb file, in presence of ADP and ATP for the 4-mers and only ATP for the 8-mers. For the ATP state, the ADP has been replaced by ATP. The N-terminal and C-terminal extremities only were generated using modeller.

For all systems, the residues have been protonated following the results of PROPKA3 [86] at pH 7.4. In our simulations, unmethylated HIS73 was found to be protonated once. The histidine was protonated in Nepsilon, as represented in the study of Wilkinson and coauthors [11] and as described by Li et al. [87]. Therefore, as the methylation has been shown to occur only in epsilon, we replaced the epsilon hydrogen by the methyl group in our simulations. We used these protonation states for all the systems presented in this study. All systems were solvated in water boxes using the xyzedit tool of the Tinker 8 distribution[88], so that there was at least 20Å between two images of the protein. The systems were then neutralized and KCL atoms were added to reach 150mM concentration. Regarding the simulations at high Mg^2+^ ions concentrations, all K+ atoms were replaced by half number of Mg^2+^ ions.

The force field parameters used for the protein parameters was the AMOEBA Polarizable force field for proteins [89]. Previously published parameters were used for the ATP and ADP systems [90]. The parameters of the HIC73 residues were developed following the procedure used to develop the AMOEBA force field for proteins. All QM calculations were performed using Gaussian09 [91]. The model residue used to develop the parameters was a dipeptide Ac-HIC-NME were the Ac, NME and backbone parameters were extracted from the AMOEBABIO18 force field [92]. Briefly, geometry optimisation were carried out at the MP2/6-31G* level. Initial atomic multipoles were derived at the MP2/6-311G** level using the Distributed Dipole Analysis (DMA) procedure [93]. The resulting atomic multipoles were optimized against MP2/aug-cc-pvtz electrostatic potential on a set of grid points distributed around the dipeptide. During this fitting, the monopoles were held fixed. The point charges were adjusted at the junction atom between the backbone and the sidechain (adjustment of 0.03 on the -CH2-carbon atom charge), to insure electrical neutrality. As realised in the original AMOEBA for protein publication, 3 conformations were used to realize the fitting of the dipole and multipole components. The valence,vdW and torsional parameters were extracted from the AMOEBA parameters of the classical HIS residue.

### Molecular Dynamics Simulation parameters

All molecular dynamics simulations were performed using the Tinker-HP software ported on GPU [94, 95]. Periodic boundary conditions were applied using the Particle Mesh Ewald method[96]. The van der Waals and PME cutoffs were respectively of 12Å and 7Å, in combination with an analytical long range correction of the vdW interactions. The dipole convergence criterion of the preconditioned conjugate gradient polarization solver were set to 0.01 Debye/atom and to 0.00001 Debye/atom for minimization and molecular dynamics simulations respectively.

During minimization, the L-BFGS optimizer has been used without polarization or electrostatics terms. The RESPA propagator (1fs timestep) and the berendsen barostat where used for equilibration steps, and the BAOAB-RESPA1 propagator [97] was used in combination with the Monte Carlo barostat for productions steps. The solvent was progressively heat up in the NVT ensemble, from 5K to 300K using 10K steps and spending 5ps at each temperature, before undergoing additional 100ps at 300K. The system was then allowed to slowly relax for 3 times 400ps in the NPT ensemble while applying harmonic restraints of 10, 5 and finally 1 kcal/mol/A on the backbone atoms of the protein. Three final equilibration steps were performed for 100ps, 200ps and 500 ps by respectively increasing the outer-timestep of the BAOAB-RESPA1 propagator from 1fs to 2fs up to 5fs.

During production, hydrogen mass repartitioning was applied on the system. A first 10ns simulations was performed to generate a first set of structures. In order to maximize the phase space exploration, we then resorted to an adaptive sampling procedure: a certain number of structures were first extracted from this initial simulation to perform the first adaptive sampling round. The seeds were then chosen following a procedure already described in [58]. Briefly, a principal component analysis is performed on the 10ns simulation using the scikit-learn [98] and MDTraj [99] packages from which the n=4 first principal modes are considered (note:10 modes are calculated). The density *ρ_k_* of the conformational space is then projected on the 4 modes and approximated using a Gaussian density kernel estimator:

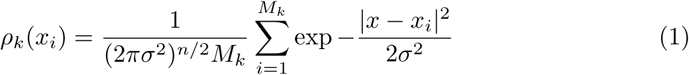

With the *σ* bandwidth being chosen with the D.W Scott method of Scipy [100], *M_k_* being the total number of configurations, *x_i_* the orthogonal projection of the configuration on the n PCA modes. The selection of new seeds *x_i_* was then biased as following :

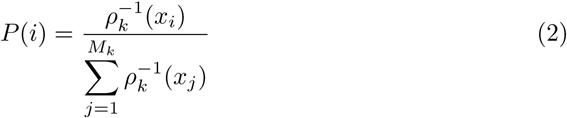

The probability of selecting the *x_i_* structure being inversely proportional to its density, projected on the first 4 PCA components, this selection methods favors the exploration of structural states

Following this, 10 ns simulations were run to form the new phase space of structures for the next adaptive sampling round. For each following round, all simulations are added to the conformational space on which the next adaptive sampling are performed. The number of seeds used for each round is summed up in Table 1.

**Table 1.** Number of seeds of each round of Adaptive sampling.

| Systems | 1st | 2nd to 10th | total simulation time |
| --- | --- | --- | --- |
| Monomers and 4-mers | 8*10ns | 16*10ns | 1.52µs |
| 8-mers | 8*10ns | 16*10ns (2 rounds) | 400ns |

**Table 2.** Definition of each subdomain.

|  | SD1 | SD2 | SD3 | SD4 |
| --- | --- | --- | --- | --- |
| Residues | 5-33<br>80-147<br>334-349 | 34-39<br>52-69 | 148-179<br>273-333 | 180-219<br>252-262 |

Regarding the 4-mer systems, we were interested in the behaviour of the barbed end only. For this extremity, it has recently been demonstrated that the behaviour of the B and B-1 residues is different than other residues of the filament [36]. Therefore, to simulate the behaviour of this extremity only, the backbone atoms of two monomers forming the pointed end have been constrained using a positional restraint of 10kcal per mol allowing these atoms to vibrate around a fixed position. For more details see p. 88 of Tinker documentation: [https://dasher.wustl.edu/tinker/downloads/tinker-guide.pdf]. This way, it was possible to study the evolution of the barbed end in a constrained filament, while keeping the sidechains free. The same strategy has been employed for 8-mer systems, constraining 2 subunits on barbed and pointed ends of the filament, to study the dynamics of the core monomers. 1.52 µs long simulations were generated on thirteen monomer systems and four 4-mer systems, and 400ns were generated on 8-mer systems, resulting in a total of 26.64 µs (see Table 1).

### Molecular Dynamics Analysis

For analysis, the data were collected each 100 ps of simulation. Measures have been performed using the VMD software [101]. The Root Mean Square Fluctuations of an amino acid i has been defined as previously described [102]:

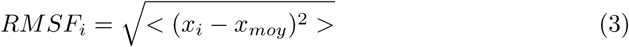

with *x_i_*the position of the center of mass of the backbone of the amino acid *i* at a certain frame, and *x_ref_* its ensemble averaged position.

Active site volumes have been computed using E-pock program [103]. We first defined a Maximum Encompassing Region (MER) by concatenating spheres to define the maximum volume of the active site as followed: a 6Å sphere around the N1, N9, PA, PG (PB for ADP) atoms of the nucleotide and one additional sphere around the Mg^2+^ ion coordinated to it. Then, for each frame of the trajectory, E-pock calculated a 3D grid based on the MER and removed grid cells which were overlapping with residues from the active site. The remaining cells defined the free volume of the active site.

Shortest paths have been calculated using the Dynamical Network Analysis (Dynetan) tool [64]. This program allowed defining nodes based on residues alpha carbons. The minimum distance between two nodes was measured on each frame of the simulation. If this distance is lower than 4.5 Å for more than 75% of the simulation, the two nodes were considered in contact. Once all nodes in contact were determined, correlation of motion was calculated between them to create the optimal correlation paths. For the calculation of allosteric paths, the shortest path of each seed has been calculated on the 10ns. Then all found shortest paths were reweighted across all seeds, and the 3 highest shortest paths have been kept for figures. All shortest paths accounting for more than 3 percent of all paths are available in supplementary information.

The dihedral and cleft angle has been defined following the work of Saunders and coworkers [29]: Dihedral = SD2-SD1-SD3-SD4, Cleft = SD2-SD4 distance. To caculate these values, the center of mass of each subdomains have been defined as displayed in table2.

Each of the observable had to be reweighted to take into account the bias introduced by the adaptive sampling. For this purpose, the unbiasing factor *α_i_* of each seed is defined as :

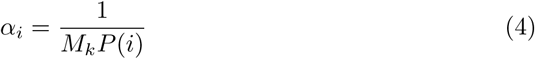

The final weight of each seed is then :

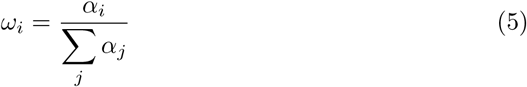

To assess whether distributions between observables were significant, Kolmogorov-Smirnov (KS) test has been performed. To apply this test, observables need to be decorrelated from each other. We took the last frame of each simulation (or seed) into account for the KS test which can be considered as independent from each other, resulting in 152 points used for each condition.

We reanalysed data from the Hummer lab available at: [https://zenodo.org/records/7765025] [104]. We have used the trajectory called replica 031 from the dataset simulations dataset 1/Metadynamics/BarbedEnd meH73p/repre-sentative trajectories corresponding to the Hummer lab enhanced sampling protocol based on simulating *swarms* of short metadynamics trajectories to open the second physically plausible pathways [105]. By methodological means, the G74-S14 backdoor was closed in the initial half of the simulation (metaD i) and then open in the final half of the simulation (metaD f).

### In vitro measurement of Pi-release rate

Human cytoplasmic *β*-actin was expressed and purified from yeast *Pichia Pastoris*, following established protocol [22]. Transformed yeast strains are a gift of Balasubramanian lab. P. pastoris transformants were revived on YPD solid media supplemented with 100 mg/L of Zeocin, and incubated overnight at 30°C. Cells were inoculated into 750 mL minimal glycerol (MGY) liquid media, and incubated overnight at 30°C, rotating at 280 rpm. Culture media was diluted by adding 6 L of fresh MGY media. Cells were cultured at 30°C, 280 rpm until optical density at 600nm reached 1.5. Cells were pelleted by centrifugation at 7,000 rpm at 25°C for 10 minutes, washed with sterilized pure water, resuspended into 6 L of minimal methanol (MM) buffer, and incubated for 24h at 30°C, 280 rpm. After 24h, 0.5% methanol was added and the culture was pursued for another 24h. Cells were pelleted by centrifugation at 7,000 rpm at 25°C for 10 minutes, washed once with water, resuspended in 30 mL pure water, and stored at −70°C as frozen beads using liquid nitrogen.

30 g of frozen cells were ground and the obtained lysate powder was resuspended into a 2X Binding buffer (20 mM imidazole, 20 mM HEPES (both pH7.4), 600 mM NaCl, 4 mM MgCl2, 2 mM ATP, 2x protease inhibitor cocktail EDTA free, 1 mM PMSF, 7 mM Beta-mercaptoethanol. Lysate was sonicated, centrifuged at 4°C, 15,000 rpm for1h, and incubated with 6 mL NTA resin at 4°C for 1 hour. The resin was pelleted down by centrifugation, washed once with 1X binding buffer, and poured into a column. Resin was then washed and equilibrated using 100 mL of G-buffer (5 mM HEPES pH 7.4, 0.2 mM CaCl2, 0.2 mM ATP, 0.5 mM DTT, and 0.01% NaN3). Chrymotrypsin was added (10 *µ*g/mL final) to the resin and incubation proceeded overnight at 4 °C. Chymotrypsin was inactivated by adding 1 mM PMSF, and lysate was incubated for 30 minutes on ice. By adding 15 mL of G-buffer, elution was collected into a single becher to recover the detached G-actin from the resin. Actin was concentrated using a Amicon 30 kDa cut-off membrane to reach 5 mL final volume. Actin was then polymerized by addition of 500 *µ*l 10x KME buffer (20 mM MgCl2, 50 mM EGTA, 1 M KCl), for 2 hours at room temperature. F-actin was pelleted by ultracentrifugation for 45 minutes, 90,000 rpm, at room temperature. F-actin pellet was re-suspended into 6 mL G-buffer, and let to depolymerize by dialysis against 1 L G-buffer at 4°C for 48 hours, with one dialysis buffer exchanged after 24h. Dialized G-actin was ultracentrifugated as above to remove any remaining F-actin. Supernatant was finally gel filtered by size exlcusion chromatography, and eluted monomeric G-actin concentration was measured by at 290 nm (*ɛ*_290_ = 0.63 mg*^−^*^1^*.mL^−^*^1^*.cm^−^*^1^).

In vitro fluorescence microscopy experiments were performed in microfluidic chambers, as described in [106]. Briefly, tens of single actin filaments were polymerized for 10-15 min from surface-anchored spectrin-actin seeds with a solution containing 1 µM monomeric *β*-actin, 1 µM human profilin and 0.5-1 µM ATP-ATTO [107] in F-buffer (10 mM Tris HCl pH 7.4, 50 mM KCl, 1 mM MgCL2, 0.2 mM EGTA, 0.2 mM ATP, 10 mM DTT, 1 mM DABCO) supplemented with 50 mM KPO4 buffer pH 7.4 to maintain actin in an ADP-Pi state. Filaments were then exposed from time t=0 to F-buffer only (without KPO4), triggering their barbed-end depolymerization and Pi release from actin subunits. Images were acquired on a Nikon Ti microscope, 60x objective, X-cite Exact lamp and Orca Flash 4.0 camera, at 1 frame every 2 to 5 s.

Movies were analyzed with a custom-made Python algorithm which tracks the depolymerizing barbed end of each filament (used packages: numpy, scipy, matplotlib, trackpy). For each filament, the spontaneous depolymerization rate was measured from the linear fit of the barbed-end position over 5 frames. The data of at least 30 filaments were pooled and the depolymerization rate over time was fitted (scipy function “curve-fit”) as [71]:

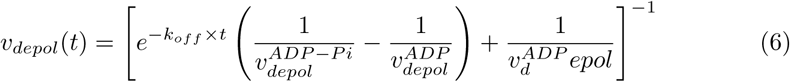

with free parameters *k_off_* the Pi release-rate, 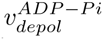 and 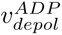 the barbed-end depolymerization rates of ADP-Pi- and ADP-actin, respectively.

## Data availability

Unless otherwise stated, all data supporting the results of this study can be found in the article, supplementary, and source data files.Data deposition: MD simulation material is available at [https://doi.org/10.5281/zenodo.14062352]. Source data are provided with this paper.

## Code availability

MD simulations were performed using the Tinker-HP v1.2: [https://github.com/TinkerTools/tinker-hp]. Scripts for visualisation are available at [https://doi.org/10.5281/zenodo.14062352]

## Acknowledgements

A. D. is funded by the ITMO Cancer of Aviesan “Mathematiques et Informatique”. This work has also received funding from the European Research Council (ERC) under the European Union’s Horizon 2020 research and innovation program (grant agreement No 810367), project EMC2 (JPP). This work was granted access to the HPC resources of CALMIP supercomputing center (under the allocation 2021-P21020) and TGCC Joliot-Curie supercomputer (under the GENCI allocation A0110712941). The Romet/Jegou lab would like to thank Mohan K. Balasubramanian for sharing the P. pastoris yeast strains to express human recombinant actin.

## Author contribution

**Designed research:** A. S., M. C., A. J., J.-P. P. ; **Performed research:** A. S., L. L., H. W., M. B. ; **Contributed analytic tools:** B. W., P. R., L. L., J.-P. P.; **Analyzed data:** A. S., H. W. ; **Wrote the paper:** A. S., A. J., M. C., J.-P. P.

## References

[1] Svitkina, T.: The Actin Cytoskeleton and Actin-Based Motility. Cold Spring Harbor Perspectives in Biology 10(1), 018267 (2018) 10.1101/cshperspect.a018267

[2] Perrin, B.J., Ervasti, J.M.: The actin gene family: Function follows isoform. Cytoskeleton 67(10), 630–634 (2010) 10.1002/cm.20475

[3] Varland, S., Vandekerckhove, J., Drazic, A.: Actin Post-translational Modifications: The Cinderella of Cytoskeletal Control. Trends in Biochemical Sciences 44(6), 502–516 (2019) 10.1016/j.tibs.2018.11.010

[4] Merino, F., Pospich, S., Raunser, S.: Towards a structural understanding of the remodeling of the actin cytoskeleton. Seminars in Cell & Developmental Biology 102, 51–64 (2020) 10.1016/j.semcdb.2019.11.018

[5] Wegner, A.: Treadmilling of actin at physiological salt concentrations: An analysis of the critical concentrations of actin filaments. Journal of Molecular Biology 161(4), 607–615 (1982) 10.1016/0022-2836(82)90411-9

[6] Oda, T., Iwasa, M., Aihara, T., Maéda, Y., Narita, A.: The nature of the globular- to fibrous-actin transition. Nature 457(7228), 441–445 (2009) 10.1038/nature07685

[7] Blanchoin, L., Pollard, T.D.: Hydrolysis of atp by polymerized actin depends on the bound divalent cation but not profilin. Biochemistry 41(2), 597–602 (2002) 10.1021/bi011214b

[8] Oosterheert, W., Blanc, F.E.C., Roy, A., Belyy, A., Sanders, M.B., Hofnagel, O., Hummer, G., Bieling, P., Raunser, S.: Molecular mechanisms of inorganic-phosphate release from the core and barbed end of actin filaments. Nature Structural & Molecular Biology 30 (2023) 10.1038/s41594-023-01101-9

[9] Reynolds, M.J., Hachicho, C., Carl, A.G., Gong, R., Alushin, G.M.: Bending forces and nucleotide state jointly regulate F-actin structure. Nature 611, 380–386 (2022) 10.1038/s41586-022-05366-w

[10] Kang, H., Bradley, M.J., McCullough, B.R., Pierre, A., Grintsevich, E.E., Reisler, E., Cruz, E.M.D.L.: Identification of cation-binding sites on actin that drive polymerization and modulate bending stiffness. Proceedings of the National Academy of Sciences 109(42), 16923–16927 (2012) 10.1073/pnas.1211078109 https://www.pnas.org/doi/pdf/10.1073/pnas.1211078109

[11] Wilkinson, A.W., Diep, J., Dai, S., Liu, S., Ooi, Y.S., Song, D., Li, T.M., Horton, J.R., Zhang, X., Liu, C., Trivedi, D.V., Ruppel, K.M., Vilches-Moure, J.G., Casey, K.M., Mak, J., Cowan, T., Elias, J.E., Nagamine, C.M., Spudich, J.A., Cheng, X., Carette, J.E., Gozani, O.: SETD3 is an actin histidine methyltransferase that prevents primary dystocia. Nature 565(7739), 372–376 (2019) 10.1038/s41586-018-0821-8

[12] Johnson, P., Harris, C.I., Perry, S.V.: 3-Methylhistidine in actin and other muscle proteins. Biochemical Journal 105(1), 361–370 (1967) 10.1042/bj1050361 https://portlandpress.com/biochemj/article-pdf/105/1/361/759780/bj1050361.pdf

[13] Kwiatkowski, S., Seliga, A.K., Vertommen, D., Terreri, M., Ishikawa, T., Grabowska, I., Tiebe, M., Teleman, A.A., Jagielski, A.K., Veiga-Da-Cunha, M., Drozak, J.: SETD3 protein is the actin-specific histidine N-methyltransferase. eLife 7(2016), 1–42 (2018) 10.7554/eLife.37921

[14] Graceffa, P., Dominguez, R.: Crystal structure of monomeric actin in the ATP state: Structural basis of nucleotide-dependent actin dynamics. Journal of Biological Chemistry 278(36), 34172–34180 (2003) 10.1074/jbc.M303689200

[15] Nyman, T., Schüler, H., Korenbaum, E., Schutt, C.E., Karlsson, R., Lindberg, U.: The role of meh73 in actin polymerization and atp hydrolysis. Journal of Molecular Biology 317(4), 577–589 (2002) 10.1006/jmbi.2002.5436

[16] Bunnell, T.M., Burbach, B.J., Shimizu, Y., Ervasti, J.M.: *β*-actin specifically controls cell growth, migration, and the g-actin pool. Molecular Biology of the Cell 22(21), 4047–4058 (2011) 10.1091/mbc.e11-06-0582 10.1091/mbc.e11-06-0582. PMID: 21900491

[17] Mahmood, S.R., Xie, X., Hosny El Said, N., Venit, T., Gunsalus, K.C., Percipalle, P.: *β*-actin dependent chromatin remodeling mediates compartment level changes in 3d genome architecture. Nature Communications 12(1), 5240 (2021) 10.1038/s41467-021-25596-2

[18] Dugina, V.B., Shagieva, G.S., Shakhov, A.S., Alieva, I.B.: The cytoplasmic actins in the regulation of endothelial cell function. International Journal of Molecular Sciences 22(15) (2021) 10.3390/ijms22157836

[19] Nietmann, P., Kaub, K., Suchenko, A., Stenz, S., Warnecke, C., Balasubramanian, M.K., Janshoff, A.: Cytosolic actin isoforms form networks with different rheological properties that indicate specific biological function. Nature Communications 14(1), 7989 (2023) 10.1038/s41467-023-43653-w

[20] Guo, C., Liu, S., Wang, J., Sun, M.-Z., Greenaway, F.T.: Actb in cancer. Clinica Chimica Acta 417, 39–44 (2013) 10.1016/j.cca.2012.12.012

[21] Ceron, R.H., Carman, P.J., Rebowski, G., Boczkowska, M., Heuckeroth, R.O., Dominguez, R.: A solution to the long-standing problem of actin expression and purification. Proceedings of the National Academy of Sciences 119(41), 2209150119 (2022) 10.1073/pnas.2209150119 https://www.pnas.org/doi/pdf/10.1073/pnas.2209150119

[22] Hatano, T., Sivashanmugam, L., Suchenko, A., Hussain, H., Balasubramanian, M.K.: Pick-ya actin – a method to purify actin isoforms with bespoke key post-translational modifications. Journal of Cell Science 133(2), 241406 (2020) 10.1242/jcs.241406 https://journals.biologists.com/jcs/article-pdf/133/2/jcs241406/1989673/jcs241406.pdf

[23] Vanommeslaeghe, K., Hatcher, E., Acharya, C., Kundu, S., Zhong, S., Shim, J., Darian, E., Guvench, O., Lopes, P., Vorobyov, I., Mackerell Jr., A.D.: Charmm general force field: A force field for drug-like molecules compatible with the charmm all-atom additive biological force fields. Journal of Computational Chemistry 31(4), 671–690 (2010) 10.1002/jcc.21367 https://onlinelibrary.wiley.com/doi/pdf/10.1002/jcc.21367

[24] Maier, J.A., Martinez, C., Kasavajhala, K., Wickstrom, L., Hauser, K.E., Simmerling, C.: ff14sb: Improving the accuracy of protein side chain and backbone parameters from ff99sb. Journal of Chemical Theory and Computation 11(8), 3696–3713 (2015) 10.1021/acs.jctc.5b00255 10.1021/acs.jctc.5b00255. PMID: 26574453

[25] Souza, P.C.T., Alessandri, R., Barnoud, J., Thallmair, S., Faustino, I., Grünewald, F., Patmanidis, I., Abdizadeh, H., Bruininks, B.M.H., Wassenaar, T.A., Kroon, P.C., Melcr, J., Nieto, V., Corradi, V., Khan, H.M., Domański, J., Javanainen, M., Martinez-Seara, H., Reuter, N., Best, R.B., Vattulainen, I., Monticelli, L., Periole, X., Tieleman, D.P., Vries, A.H., Marrink, S.J.: Martini 3: a general purpose force field for coarse-grained molecular dynamics. Nature Methods 18(4), 382–388 (2021) 10.1038/s41592-021-01098-3

[26] Kim, Y.C., Hummer, G.: Coarse-grained models for simulations of multiprotein complexes: Application to ubiquitin binding. Journal of Molecular Biology 375(5), 1416–1433 (2008) 10.1016/j.jmb.2007.11.063

[27] Chu, J.-W., Voth, G.A.: Allostery of actin filaments: Molecular dynamics simulations and coarse-grained analysis. Proceedings of the National Academy of Sciences 102(37), 13111–13116 (2005) 10.1073/pnas.0503732102 https://www.pnas.org/doi/pdf/10.1073/pnas.0503732102

[28] Splettstoesser, T., Noé, F., Oda, T., Smith, J.C.: Nucleotide-dependence of g-actin conformation from multiple molecular dynamics simulations and observation of a putatively polymerization-competent superclosed state. Proteins: Structure, Function, and Bioinformatics 76(2), 353–364 (2009) 10.1002/prot.22350 https://onlinelibrary.wiley.com/doi/pdf/10.1002/prot.22350

[29] Saunders, M.G., Tempkin, J., Weare, J., Dinner, A.R., Roux, B., Voth, G.A.: Nucleotide regulation of the structure and dynamics of G-actin. Biophysical Journal 106(8), 1710–1720 (2014) 10.1016/j.bpj.2014.03.012

[30] Jepsen, L., Sept, D.: Effects of nucleotide and end-dependent actin conformations on polymerization. Biophysical Journal 119(9), 1800–1810 (2020) 10.1016/j.bpj.2020.09.024

[31] Saunders, M.G., Voth, G.A.: Water molecules in the nucleotide binding cleft of actin: Effects on subunit conformation and implications for ATP hydrolysis. Journal of Molecular Biology 413(1), 279–291 (2011) 10.1016/j.jmb.2011.07.068

[32] McCullagh, M., Saunders, M.G., Voth, G.A.: Unraveling the mystery of atp hydrolysis in actin filaments. Journal of the American Chemical Society 136(37), 13053–13058 (2014) 10.1021/ja507169f 10.1021/ja507169f. PMID: 25181471

[33] Rennebaum, S., Caflisch, A.: Inhibition of interdomain motion in g-actin by the natural product latrunculin: A molecular dynamics study. Proteins: Structure, Function, and Bioinformatics 80(8), 1998–2008 (2012) 10.1002/prot.24088 https://onlinelibrary.wiley.com/doi/pdf/10.1002/prot.24088

[34] Helal, M.A., Khalifa, S., Ahmed, S.: Differential binding of latrunculins to g-actin: A molecular dynamics study. Journal of Chemical Information and Modeling 53(9), 2369–2375 (2013) 10.1021/ci400317j 10.1021/ci400317j. PMID: 23988111

[35] Splettstoesser, T., Holmes, K.C., Noé, F., Smith, J.C.: Structural modeling and molecular dynamics simulation of the actin filament. Proteins: Structure, Function, and Bioinformatics 79(7), 2033–2043 (2011) 10.1002/prot.23017 https://onlinelibrary.wiley.com/doi/pdf/10.1002/prot.23017

[36] Zsolnay, V., Katkar, H.H., Chou, S.Z., Pollard, T.D., Voth, G.A.: Structural basis for polarized elongation of actin filaments. Proceedings of the National Academy of Sciences of the United States of America 117(48), 30458–30464 (2020) 10.1073/pnas.2011128117

[37] Schroer, C.F.E., Baldauf, L., Buren, L., Wassenaar, T.A., Melo, M.N., Koenderink, G.H., Marrink, S.J.: Charge-dependent interactions of monomeric and filamentous actin with lipid bilayers. Proceedings of the National Academy of Sciences 117(11), 5861–5872 (2020) 10.1073/pnas.1914884117 https://www.pnas.org/doi/pdf/10.1073/pnas.1914884117

[38] Shamloo, A., Mehrafrooz, B.: Nanomechanics of actin filament: A molecular dynamics simulation. Cytoskeleton 75(3), 118–130 (2018) 10.1002/cm.21429 https://onlinelibrary.wiley.com/doi/pdf/10.1002/cm.21429

[39] Jaswandkar, S.V., Faisal, H.M.N., Katti, K.S., Katti, D.R.: Dissociation mechanisms of g-actin subunits govern deformation response of actin filament. Biomacromolecules 22(2), 907–917 (2021) 10.1021/acs.biomac.0c01602 10.1021/acs.biomac.0c01602. PMID: 33481563

[40] Horan, B.G., Hall, A.R., Vavylonis, D.: Insights into actin polymerization and nucleation using a coarse-grained model. Biophysical Journal 119(3), 553–566 (2020) 10.1016/j.bpj.2020.06.019

[41] Castaneda, N., Lee, M., Rivera-Jacquez, H.J., Marracino, R.R., Merlino, T.R., Kang, H.: Actin filament mechanics and structure in crowded environments. The Journal of Physical Chemistry B 123(13), 2770–2779 (2019) 10.1021/acs.jpcb.8b12320 10.1021/acs.jpcb.8b12320. PMID: 30817154

[42] Song, C., Weichbrodt, C., Salnikov, E.S., Dynowski, M., Forsberg, B.O., Bechinger, B., Steinem, C., Groot, B.L., Zachariae, U., Zeth, K.: Crystal structure and functional mechanism of a human antimicrobial membrane channel. Proceedings of the National Academy of Sciences 110(12), 4586–4591 (2013) 10.1073/pnas.1214739110 https://www.pnas.org/doi/pdf/10.1073/pnas.1214739110

[43] Kopec, W., Köpfer, D.A., Vickery, O.N., Bondarenko, A.S., Jansen, T.L.C., Groot, B.L., Zachariae, U.: Direct knock-on of desolvated ions governs strict ion selectivity in k+ channels. Nature Chemistry 10(8), 813–820 (2018) 10.1038/s41557-018-0105-9

[44] Bellissent-Funel, M.-C., Hassanali, A., Havenith, M., Henchman, R., Pohl, P., Sterpone, F., Spoel, D., Xu, Y., Garcia, A.E.: Water determines the structure and dynamics of proteins. Chemical Reviews 116(13), 7673–7697 (2016) 10.1021/acs.chemrev.5b00664 10.1021/acs.chemrev.5b00664. PMID: 27186992

[45] Hocky, G.M., Baker, J.L., Bradley, M.J., Sinitskiy, A.V., De La Cruz, E.M., Voth, G.A.: Cations stiffen actin filaments by adhering a key structural element to adjacent subunits. The Journal of Physical Chemistry B 120(20), 4558–4567 (2016) 10.1021/acs.jpcb.6b02741 10.1021/acs.jpcb.6b02741. PMID: 27146246

[46] Kadaoluwa Pathirannahalage, S.P., Meftahi, N., Elbourne, A., Weiss, A.C.G., McConville, C.F., Padua, A., Winkler, D.A., Costa Gomes, M., Greaves, T.L., Le, T.C., Besford, Q.A., Christofferson, A.J.: Systematic comparison of the structural and dynamic properties of commonly used water models for molecular dynamics simulations. Journal of Chemical Information and Modeling 61(9), 4521–4536 (2021) 10.1021/acs.jcim.1c00794 10.1021/acs.jcim.1c00794. PMID: 34406000

[47] Shi, Y., Ren, P., Schnieders, M., Piquemal, J.-P.: 2. Polarizable Force Fields for Biomolecular Modeling, pp. 51–86. John Wiley & Sons, Ltd,(2015). 10.1002/9781118889886.ch2. https://onlinelibrary.wiley.com/doi/abs/10.1002/9781118889886.ch2

[48] Melcr, J., Piquemal, J.-P.: Accurate biomolecular simulations account for electronic polarization. Frontiers in Molecular Biosciences 6 (2019) 10.3389/fmolb.2019.00143

[49] Jing, Z., Liu, C., Cheng, S.Y., Qi, R., Walker, B.D., Piquemal, J.-P., Ren, P.: Polarizable force fields for biomolecular simulations: Recent advances and applications. Annual Review of Biophysics 48(1), 371–394 (2019) 10.1146/annurev-biophys-070317-033349 10.1146/annurev-biophys-070317-033349. PMID: 30916997

[50] Jing, Z., Rackers, J.A., Pratt, L.R., Liu, C., Rempe, S.B., Ren, P.: Thermodynamics of ion binding and occupancy in potassium channels. Chem. Sci. 12, 8920–8930 (2021) 10.1039/D1SC01887F

[51] Lynch, C.I., Klesse, G., Rao, S., Tucker, S.J., Sansom, M.S.P.: Water nanoconfined in a hydrophobic pore: Molecular dynamics simulations of transmembrane protein 175 and the influence of water models. ACS Nano 15(12), 19098–19108 (2021) 10.1021/acsnano.1c06443 10.1021/acsnano.1c06443. PMID: 34784172

[52] El Ahdab, D., Lagardère, L., Inizan, T.J., Célerse, F., Liu, C., Adjoua, O., Jolly, L.-H., Gresh, N., Hobaika, Z., Ren, P., Maroun, R.G., Piquemal, J.-P.: Interfacial water many-body effects drive structural dynamics and allosteric interactions in sars-cov-2 main protease dimerization interface. The Journal of Physical Chemistry Letters 12(26), 6218–6226 (2021) 10.1021/acs.jpclett.1c01460. PMID: 34196568

[53] Célerse, F., Lagardère, L., Derat, E., Piquemal, J.-P.: Massively parallel implementation of steered molecular dynamics in tinker-hp: Comparisons of polarizable and non-polarizable simulations of realistic systems. Journal of Chemical Theory and Computation 15(6), 3694–3709 (2019) 10.1021/acs.jctc.9b00199 10.1021/acs.jctc.9b00199. PMID: 31059250

[54] Kratochvil, H.T., Carr, J.K., Matulef, K., Annen, A.W., Li, H., Maj, M., Ostmeyer, J., Serrano, A.L., Raghuraman, H., Moran, S.D., Skinner, J.L., Perozo, E., Roux, B., Valiyaveetil, F.I., Zanni, M.T.: Instantaneous ion configurations in the k¡sup¿+¡/sup¿ ion channel selectivity filter revealed by 2d ir spectroscopy. Science 353(6303), 1040–1044 (2016) 10.1126/science.aag1447 https://www.science.org/doi/pdf/10.1126/science.aag1447

[55] Li, H., Ngo, V., Silva, M.C.D., Salahub, D.R., Callahan, K., Roux, B., Noskov, S.Y.: Representation of Ion–Protein Interactions Using the Drude Polarizable Force-Field. The Journal of Physical Chemistry B 119(29), 9401–9416 (2015) 10.1021/jp510560k

[56] El Khoury, L., Jing, Z., Cuzzolin, A., Deplano, A., Loco, D., Sattarov, B., Hédin, F., Wendeborn, S., Ho, C., El Ahdab, D., Jaffrelot Inizan, T., Sturlese, M., Sosic, A., Volpiana, M., Lugato, A., Barone, M., Gatto, B., Macchia, M.L., Bellanda, M., Battistutta, R., Salata, C., Kondratov, I., Iminov, R., Khairulin, A., Mykhalonok, Y., Pochepko, A., Chashka-Ratushnyi, V., Kos, I., Moro, S., Montes, M., Ren, P., Ponder, J.W., Lagardère, L., Piquemal, J.-P., Sabbadin, D.: Computationally driven discovery of sars-cov-2 mpro inhibitors: from design to experimental validation. Chem. Sci. 13, 3674–3687 (2022) 10.1039/D1SC05892D

[57] Blazhynska, M., Lagardère, L., Liu, C., Adjoua, O., Ren, P., Piquemal, J.-P.: Water–glycan interactions drive the sars-cov-2 spike dynamics: insights into glycan-gate control and camouflage mechanisms. Chem. Sci. 15, 14177–14187 (2024) 10.1039/D4SC04364B

[58] Jaffrelot Inizan, T., Célerse, F., Adjoua, O., El Ahdab, D., Jolly, L.-H., Liu, C., Ren, P., Montes, M., Lagarde, N., Lagardère, L., Monmarché, P., Piquemal, J.-P.: High-resolution mining of the SARS-CoV-2 main protease conformational space: supercomputer-driven unsupervised adaptive sampling. Chemical Science 12(2003) (2021) 10.1039/d1sc00145k

[59] Bowman, G.R., Ensign, D.L., Pande, V.S.: Enhanced modeling via network theory: Adaptive sampling of markov state models Journal of Chemical The-ory and Computation 6(3), 787–794 (2010) 10.1021/ct900620b 10.1021/ct900620b. PMID: 23626502

[60] Miao, Y., Feher, V.A., McCammon, J.A.: Gaussian accelerated molecular dynamics: Unconstrained enhanced sampling and free energy calculation. Journal of Chemical Theory and Computation 11(8), 3584–3595 (2015) 10.1021/acs.jctc.5b00436 10.1021/acs.jctc.5b00436. PMID: 26300708

[61] Célerse, F., Inizan, T.J., Lagardère, L., Adjoua, O., Monmarché, P., Miao, Y., Derat, E., Piquemal, J.-P.: An efficient gaussian-accelerated molecular dynamics (gamd) multilevel enhanced sampling strategy: Application to polarizable force fields simulations of large biological systems. Journal of Chemical Theory and Computation 18(2), 968–977 (2022) 10.1021/acs.jctc.1c01024. PMID: 35080892

[62] Ma, W., You, S., Regnier, M., McCammon, J.A.: Integrating comparative modeling and accelerated simulations reveals conformational and energetic basis of actomyosin force generation. Proceedings of the National Academy of Sciences 120(9), 2215836120 (2023) 10.1073/pnas.2215836120 https://www.pnas.org/doi/pdf/10.1073/pnas.2215836120

[63] Westerlund, A.M., Fleetwood, O., Pérez-Conesa, S., Delemotte, L.: Network analysis reveals how lipids and other cofactors influence membrane protein allostery. J. Chem. Phys. 153(14), 141103 (2020)

[64] Melo, M.C.R., Bernardi, R.C., De La Fuente-Nunez, C., Luthey-Schulten, Z.: Generalized correlation-based dynamical network analysis: A new high-performance approach for identifying allosteric communications in molecular dynamics trajectories. Journal of Chemical Physics 153(13) (2020) 10.1063/5.0018980

[65] Rubenstein, P.A., Wen, K.-K.: Insights into the effects of disease-causing mutations in human actins. Cytoskeleton 71(4), 211–229 (2014) 10.1002/cm.21169 https://onlinelibrary.wiley.com/doi/pdf/10.1002/cm.21169

[66] Ali, R., Zahm, J.A., Rosen, M.K.: Bound nucleotide can control the dynamic architecture of monomeric actin. Nature Structural & Molecular Biology 29(4), 320–328 (2022) 10.1038/s41594-022-00743-5

[67] Kruth, K.A., Rubenstein, P.A.: Two deafness-causing (dfna20/26) actin mutations affect arp2/3-dependent actin regulation*. Journal of Biological Chemistry 287(32), 27217–27226 (2012) 10.1074/jbc.M112.377283

[68] Carman, P.J., Barrie, K.R., Rebowski, G., Dominguez, R.: Structures of the free and capped ends of the actin filament. Science 380(6651), 1287–1292 (2023) 10.1126/science.adg6812 https://www.science.org/doi/pdf/10.1126/science.adg6812

[69] Chou, S.Z., Pollard, T.D.: Mechanism of actin polymerization revealed by cryo-EM structures of actin filaments with three different bound nucleotides. Proceedings of the National Academy of Sciences of the United States of America 116(10), 4265–4274 (2019) 10.1073/pnas.1807028115

[70] Wang, Y., Wu, J., Zsolnay, V., Pollard, T.D., Voth, G.A.: Mechanism of phosphate release from actin filaments. Proceedings of the National Academy of Sciences 121(29), 2408156121 (2024) 10.1073/pnas.2408156121 https://www.pnas.org/doi/pdf/10.1073/pnas.2408156121

[71] Jégou, A., Niedermayer, T., Orbán, J., Didry, D., Lipowsky, R., Carlier, M.-F., Romet-Lemonne, G.: Individual actin filaments in a microfluidic flow reveal the mechanism of atp hydrolysis and give insight into the properties of profilin. PLOS Biology 9(9), 1–10 (2011) 10.1371/journal.pbio.1001161

[72] Carloni, P., Sanbonmatsu, K.: Exascale simulations and beyond. Current Opinion in Structural Biology 89, 102939 (2024) 10.1016/j.sbi.2024.102939

[73] Guo, Q., Liao, S., Kwiatkowski, S., Tomaka, W., Yu, H., Wu, G., Tu, X., Min, J., Drozak, J., Xu, C.: Structural insights into SETD3-mediated histidine methylation on *β*-actin. Elife 8 (2019)

[74] De La Cruz, E.M., Roland, J., McCullough, B.R., Blanchoin, L., Martiel, J.-L.: Origin of twist-bend coupling in actin filaments. Biophys. J. 99(6), 1852–1860 (2010)

[75] Cao, W., Goodarzi, J.P., De La Cruz, E.M.: Energetics and kinetics of cooperative cofilin-actin filament interactions. J. Mol. Biol. 361(2), 257–267 (2006)

[76] Wioland, H., Guichard, B., Senju, Y., Myram, S., Lappalainen, P., Jégou, A., Romet-Lemonne, G.: ADF/Cofilin accelerates actin dynamics by severing filaments and promoting their depolymerization at both ends. Curr. Biol. 27(13), 1956–19677 (2017)

[77] Wioland, H., Jegou, A., Romet-Lemonne, G.: Torsional stress generated by adf/cofilin on cross-linked actin filaments boosts their severing. Proceedings of the National Academy of Sciences 116(7), 2595–2602 (2019) 10.1073/pnas.1812053116 https://www.pnas.org/doi/pdf/10.1073/pnas.1812053116

[78] Lappalainen, P., Kotila, T., Jégou, A., Romet-Lemonne, G.: Biochemical and mechanical regulation of actin dynamics. Nat. Rev. Mol. Cell Biol. 23 (2022)

[79] Kanematsu, Y., Narita, A., Oda, T., Koike, R., Ota, M., Takano, Y., Moritsugu, K., Fujiwara, I., Tanaka, K., Komatsu, H., Nagae, T., Watanabe, N., Iwasa, M., Maéda, Y., Takeda, S.: Structures and mechanisms of actin ATP hydrolysis. Proc. Natl. Acad. Sci. U. S. A. 119(43), 2122641119 (2022)

[80] Kwiatkowski, S., Drozak, J.: Protein histidine methylation. Curr Protein Pept Sci 21(7), 675–689 (2020)

[81] Kapell, S., Jakobsson, M.E.: Large-scale identification of protein histidine methylation in human cells. NAR Genomics and Bioinformatics 3(2), 045 (2021) 10.1093/nargab/lqab045 https://academic.oup.com/nargab/article-pdf/3/2/lqab045/53692815/lqab045.pdf

[82] Otterbein, L.R., Graceffa, P., Dominguez, R.: The crystal structure of uncomplexed actin in the ADP state. Science 293(5530), 708–711 (2001) https://doi. org/10.1126/science.1059700

[83] Chen, L., Kashina, A.: Post-translational modifications of the protein termini. Frontiers in Cell and Developmental Biology 9 (2021) 10.3389/fcell.2021.719590

[84] Šali, A., Blundell, T.L.: Comparative Protein Modelling by Satisfaction of Spatial Restraints (1993)

[85] Gurel, P.S., Kim, L.Y., Ruijgrok, P.V., Omabegho, T., Bryant, Z., Alushin, G.M.: Cryo-EM structures reveal specialization at the myosin VI-actin interface and a mechanism of force sensitivity. eLife 6(Md), 1–33 (2017) 10.7554/eLife.31125

[86] Olsson, M.H.M., SØndergaard, C.R., Rostkowski, M., Jensen, J.H.: PROPKA3: Consistent treatment of internal and surface residues in empirical p K a predictions. Journal of Chemical Theory and Computation 7(2), 525–537 (2011) 10.1021/ct100578z

[87] Li, S., Hong, M.: Protonation, tautomerization, and rotameric structure of histidine: A comprehensive study by magic-angle-spinning solid-state nmr. Journal of the American Chemical Society 133(5), 1534–1544 (2011) 10.1021/ja108943n 10.1021/ja108943n. PMID: 21207964

[88] Rackers, J.A., Wang, Z., Lu, C., Laury, M.L., Lagardère, L., Schnieders, M.J., Piquemal, J.P., Ren, P., Ponder, J.W.: Tinker 8: Software Tools for Molecular Design. Journal of Chemical Theory and Computation 14(10), 5273–5289 (2018) 10.1021/acs.jctc.8b00529

[89] Shi, Y., Xia, Z., Zhang, J., Best, R., Wu, C., Ponder, J.W., Ren, P.: Polarizable atomic multipole-based AMOEBA force field for proteins. Journal of Chemical Theory and Computation 9(9), 4046–4063 (2013) 10.1021/ct4003702

[90] Walker, B., Jing, Z., Ren, P.: Molecular dynamics free energy simulations of ATP:Mg2+ and ADP:Mg2+ using the polarisable force field AMOEBA. Molecular Simulation 0(0), 1–10 (2020) 10.1080/08927022.2020.1725003

[91] Frisch, M.J., Trucks, G.W., Schlegel, H.B., Scuseria, G.E., Robb, M.A., Cheeseman, J.R., Scalmani, G., Barone, V., Mennucci, B., Petersson, G.A., Nakatsuji, H., Caricato, M., Li, X., Hratchian, H.P., Izmaylov, A.F., Bloino, J., Zheng, G., Sonnenberg, J.L., Hada, M., Ehara, M., Toyota, K., Fukuda, R., Hasegawa, J., Ishida, M., Nakajima, T., Honda, Y., Kitao, O., Nakai, H., Vreven, T., Montgomery, J.A., Peralta, J.E., Ogliaro, F., Bearpark, M., Heyd, J.J., Brothers, E., Kudin, K.N., Staroverov, V.N., Kobayashi, R., Normand, J., Raghavachari, K., Rendell, A., Burant, J.C., Iyengar, S.S., Tomasi, J., Cossi, M., Rega, N., Millam, J.M., Klene, M., Knox, J.E., Cross, J.B., Bakken, V., Adamo, C., Jaramillo, J., Gomperts, R., Stratmann, R.E., Yazyev, O., Austin, A.J., Cammi, R., Pomelli, C., Ochterski, J.W., Martin, R.L., Morokuma, K., Zakrzewski, V.G., Voth, G.A., Salvador, P., Dannenberg, J.J., Dapprich, S., Daniels, A.D., Farkas, Ö., Foresman, J.B., Ortiz, J.V., Cioslowski, J., Fox, D.J.: Gaussian 09 Revision A.2 (2009)

[92] Zhang, C., Lu, C., Jing, Z., Wu, C., Piquemal, J.-P., Ponder, J.W., Ren, P.: Amoeba polarizable atomic multipole force field for nucleic acids. Journal of Chemical Theory and Computation 14(4), 2084–2108 (2018) 10.1021/acs.jctc.7b01169 10.1021/acs.jctc.7b01169. PMID: 29438622

[93] Stone, A.J.: Distributed multipole analysis, or how to describe a molecular charge distribution. Chemical Physics Letters 83(2), 233–239 (1981) 10.1016/0009-2614(81)85452-8

[94] Lagardère, L., Jolly, L.-H., Lipparini, F., Aviat, F., Stamm, B., Jing, Z.F., Harger, M., Torabifard, H., Cisneros, G.A., Schnieders, M.J., Gresh, N., Maday, Y., Ren, P.Y., Ponder, J.W., Piquemal, J.-P.: Tinker-hp: a massively parallel molecular dynamics package for multiscale simulations of large complex systems with advanced point dipole polarizable force fields. Chem. Sci. 9, 956–972 (2018) 10.1039/C7SC04531J

[95] Adjoua, O., Lagardère, L., Jolly, L.H., Durocher, A., Very, T., Dupays, I., Wang, Z., Inizan, T.J., Célerse, F., Ren, P., Ponder, J.W., Piquemal, J.P.: Tinker-HP: Accelerating Molecular Dynamics Simulations of Large Complex Systems with Advanced Point Dipole Polarizable Force Fields Using GPUs and Multi-GPU Systems. Journal of Chemical Theory and Computation 17(4), 2034–2053 (2021) 10.1021/acs.jctc.0c01164 [arXiv:2011.01207](https://arxiv.org/abs/2011.01207)

[96] Lagardère, L., Lipparini, F., Polack, É., Stamm, B., Cancès, É., Schnieders, M., Ren, P., Maday, Y., Piquemal, J.P.: Scalable Evaluation of Polarization Energy and Associated Forces in Polarizable Molecular Dynamics: II. Toward Massively Parallel Computations Using Smooth Particle Mesh Ewald. Journal of Chemical Theory and Computation 11(6), 2589–2599 (2015) 10.1021/acs.jctc.5b00171

[97] Lagardère, L., Aviat, F., Piquemal, J.P.: Pushing the Limits of Multiple-Time-Step Strategies for Polarizable Point Dipole Molecular Dynamics. Journal of Physical Chemistry Letters 10(10), 2593–2599 (2019) 10.1021/acs.jpclett.9b00901

[98] Pedregosa, F., Varoquaux, G., Gramfort, A., Michel, V., Thirion, B., Grisel, O., Blondel, M., Prettenhofer, P., Weiss, R., Dubourg, V., Vanderplas, J., Passos, A., Cournapeau, D., Brucher, M., Perrot, M., Duchesnay, E.: Scikit-learn: Machine Learning in Python. JMLR 12(82), 2825–2830 (2011)

[99] McGibbon, R.T., Beauchamp, K.A., Harrigan, M.P., Klein, C., Swails, J.M., Herńandez, C.X., Schwantes, C.R., Wang, L.P., Lane, T.J., Pande, V.S.: MDTraj: A Modern Open Library for the Analysis of Molecular Dynamics Trajectories. Biophysical Journal 109(8), 1528–1532 (2015) 10.1016/j.bpj.2015.08.015

[100] Virtanen, P., Gommers, R., Oliphant, T.E., Haberland, M., Reddy, T., Cournapeau, D., Burovski, E., Peterson, P., Weckesser, W., Bright, J., Walt, S.J., Brett, M., Wilson, J., Millman, K.J., Mayorov, N., Nelson, A.R.J., Jones, E., Kern, R., Larson, E., Carey, C.J., Polat, I., Feng, Y., Moore, E.W., VanderPlas, J., Laxalde, D., Perktold, J., Cimrman, R., Henriksen, I., Quintero, E.A., Harris, C.R., Archibald, A.M., Ribeiro, A.H., Pedregosa, F., Mulbregt, P., Vijaykumar, A., Bardelli, A.P., Rothberg, A., Hilboll, A., Kloeckner, A., Scopatz, A., Lee, A., Rokem, A., Woods, C.N., Fulton, C., Masson, C., Häggström, C., Fitzgerald, C., Nicholson, D.A., Hagen, D.R., Pasechnik, D.V., Olivetti, E., Martin, E., Wieser, E., Silva, F., Lenders, F., Wilhelm, F., Young, G., Price, G.A., Ingold, G.L., Allen, G.E., Lee, G.R., Audren, H., Probst, I., Dietrich, J.P., Silterra, J., Webber, J.T., Slavič, J., Nothman, J., Buchner, J., Kulick, J., Schönberger, J.L., de Miranda Cardoso, J.V., Reimer, J., Harrington, J., Rodríguez, J.L.C., Nunez-Iglesias, J., Kuczynski, J., Tritz, K., Thoma, M., Newville, M., Kümmerer, M., Bolingbroke, M., Tartre, M., Pak, M., Smith, N.J., Nowaczyk, N., Shebanov, N., Pavlyk, O., Brodtkorb, P.A., Lee, P., McGibbon, R.T., Feldbauer, R., Lewis, S., Tygier, S., Sievert, S., Vigna, S., Peterson, S., More, S., Pudlik, T., Oshima, T., Pingel, T.J., Robitaille, T.P., Spura, T., Jones, T.R., Cera, T., Leslie, T., Zito, T., Krauss, T., Upadhyay, U., Halchenko, Y.O., Vázquez-Baeza, Y.: SciPy 1.0: fundamental algorithms for scientific computing in Python. Nature Methods 17(3), 261–272 (2020) 10.1038/s41592-019-0686-2 [arXiv:1907.10121](https://arxiv.org/abs/1907.10121)

[101] Humphrey, W., Dalke, A., Schulten, K.: VMD: Visual Molecular Dynamics. Journal of molecular graphics 14(October 1995), 33–38 (1996) [arXiv:arXiv:1503.05249v1](https://arxiv.org/abs/arXiv:1503.05249v1)

[102] Welford, B.P.: Note on a method for calculating corrected sums of squares and products. Technometrics 4(3), 419–420 (1962) 10.1080/00401706.1962.10490022 https://www.tandfonline.com/doi/pdf/10.1080/00401706.1962.10490022

[103] Laurent, B., Chavent, M., Cragnolini, T., Dahl, A.C.E., Pasquali, S., Derreumaux, P., Sansom, M.S.P., Baaden, M.: Epock: Rapid analysis of protein pocket dynamics. Bioinformatics 31(9), 1478–1480 (2015) 10.1093/bioinformatics/btu822

[104] Blanc, F.E.C.: MD Simulations Molecular mechanisms of inorganic-phosphate release from the core and barbed end of actin filaments, 10.5281/zenodo.7765025. Zenodo (2023). 10.5281/zenodo.7765025. 10.5281/zenodo.7765025

[105] Okazaki, K.-i., Hummer, G.: Phosphate release coupled to rotary motion of F1-ATPase. Proceedings of the National Academy of Sciences 110(41), 16468–16473 (2013) 10.1073/pnas.1305497110

[106] AU - Wioland, H., AU - Ghasemi, F., AU - Chikireddy, J., AU - Romet-Lemonne, G., AU - Jégou, A.: Using microfluidics and fluorescence microscopy to study the assembly dynamics of single actin filaments and bundles. JoVE 183(183), 63891 (2022) 10.3791/63891

[107] Colombo, J., Antkowiak, A., Kogan, K., Kotila, T., Elliott, J., Guillotin, A., Lappalainen, P., Michelot, A.: A functional family of fluorescent nucleotide analogues to investigate actin dynamics and energetics. Nature Communications 12(1), 548 (2021) 10.1038/s41467-020-20827-4

