## Supplementary Information for "Histidine 73 methylation coordinates *β*-actin plasticity in response to key environmental factors"

### Supplementary informations for : Histidine 73 methylation coordinates $\beta$-actin plasticity in response to key environmental factors

Adrien Schahl[1,2], Louis Lagardère[2], Brandon Walker[3], Pengyu Ren[3], Hugo Wioland[4], Maya Ballet[4], Antoine Jégou[4], Matthieu Chavent[1,5*], Jean-Philip Piquemal[2*]

[1]Institut de Pharmacologie et de Biologie Structurale, Université de Toulouse, CNRS, Toulouse, 31400, France.
[2]Laboratoire de Chimie Théorique, Sorbonne Université, UMR 7616 CNRS, Paris, F-75005, France.
[3]Department of Biomedical Engineering, The University of Texas at Austin, Austin, 78712, Texas, United States.
[4]Institut Jacques Monod, Université Paris Cité, CNRS, , Paris, F-75013, France.
[5]Laboratoire de Microbiologie et Génétique Moléculaires (LMGM), Centre de Biologie Intégrative (CBI), Université de Toulouse, CNRS, , Toulouse, 31400, France.

Contributing authors:;

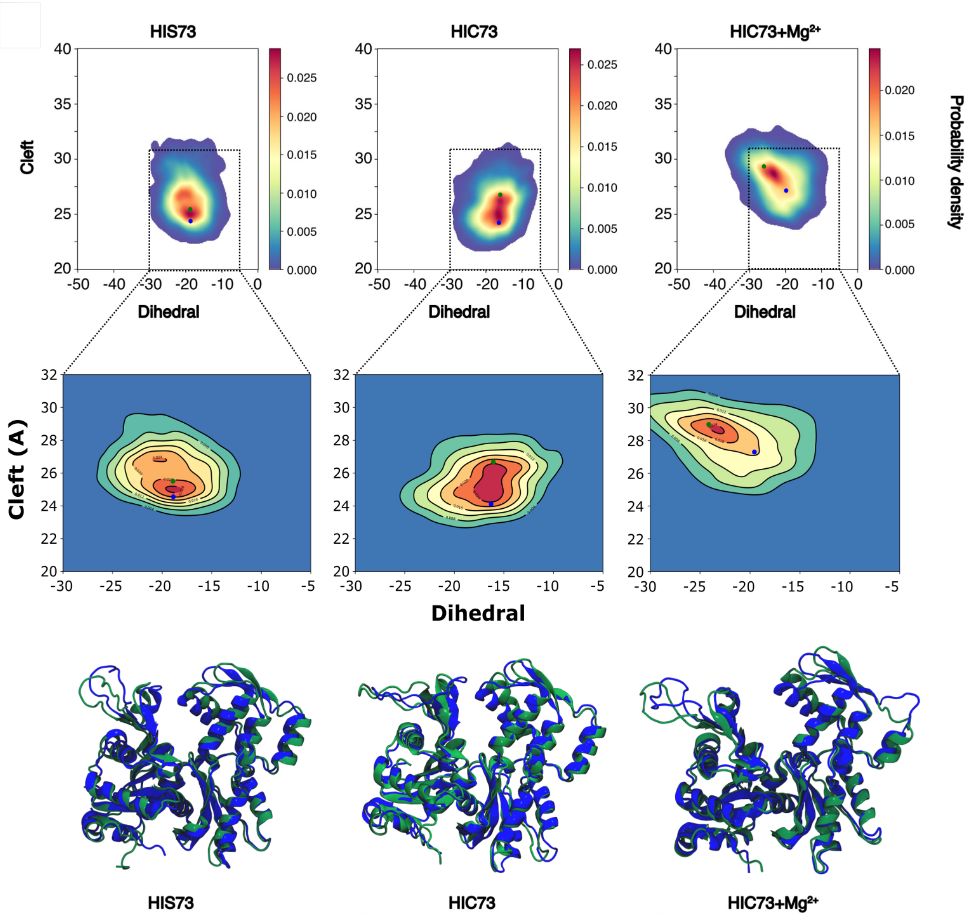


**Supplementary Figure 1 Zoomed contourplot of Cleft-Diedral maps referenced in figure 1, panel B for non-methylated and methylated $\beta$-actin in the ATP state**. In the bottom row are superpositions of structures corresponding to the green and blue dots projections on the maps. Supplementary movies 1,2 and 3 are showing morphings of these conformations for simulations of HIS, HIC and HIC + $Mg^{2+}$ respectively.

```
Alpha actin        1 DEDETTALVCDNGSGLVKAGFAGDDAPRAVFPSIVGRPRHQGVMVGMGQK     50
                     .:|:..|||.|||||:.|||||||||||||||||||||||||||||||||
Beta actin         1 MDDDIAALVVDNGSGMCKAGFAGDDAPRAVFPSIVGRPRHQGVMVGMGQK     50

Alpha actin       51 DSYVGDEAQSKRGILTLKYPIEHGIITNWDDMEKIWHHTFYNELRVAPEE    100
                     |||||||||||||||||||||||||:||||||||||||||||||||||||
Beta actin        51 DSYVGDEAQSKRGILTLKYPIEHGIVTNWDDMEKIWHHTFYNELRVAPEE    100

Alpha actin      101 HPTLLTEAPLNPKANREKMTQIMFETFNVPAMYVAIQAVLSLYASGRTTG    150
                     ||.|||||||||||||||||||||||||.|||||||||||||||||||||
Beta actin       101 HPVLLTEAPLNPKANREKMTQIMFETFNTPAMYVAIQAVLSLYASGRTTG    150

Alpha actin      151 IVLDSGDGVTHNVPIYEGYALPHAIMRLDLAGRDLTDYLMKILTERGYSF    200
                     ||:||||||||.|||||||||||||:||||||||||||||||||||||||
Beta actin       151 IVMDSGDGVTHTVPIYEGYALPHAILRLDLAGRDLTDYLMKILTERGYSF    200

Alpha actin      201 VTTAEREIVRDIKEKLCYVALDFENEMATAASSSSLEKSYELPDGQVITI    250
                     .|||||||||||||||||||||||.|||||||||||||||||||||||||
Beta actin       201 TTTAEREIVRDIKEKLCYVALDFEQEMATAASSSSLEKSYELPDGQVITI    250

Alpha actin      251 GNERFRCPETLFQPSFIGMESAGIHETTYNSIMKCDIDIRKDLYANNVMS    300
                     |||||||||.||||||:||||.||||||:|||||||:|||||||||.|:|
Beta actin       251 GNERFRCPEALFQPSFLGMESCGIHETTFNSIMKCDVDIRKDLYANTVLS    300

Alpha actin      301 GGTTMYPGIADRMQKEITALAPSTMKIKIIAPPERKYSVWIGGSILASLS    350
                     ||||||||||||||||||||||||||||||||||||||||||||||||||
Beta actin       301 GGTTMYPGIADRMQKEITALAPSTMKIKIIAPPERKYSVWIGGSILASLS    350

Alpha actin      351 TFQQMWITKQEYDEAGPSIVHRKCF    375
                     |||||||:||||||:||||||||||
Beta actin       351 TFQQMWISKQEYDESGPSIVHRKCF    375
```

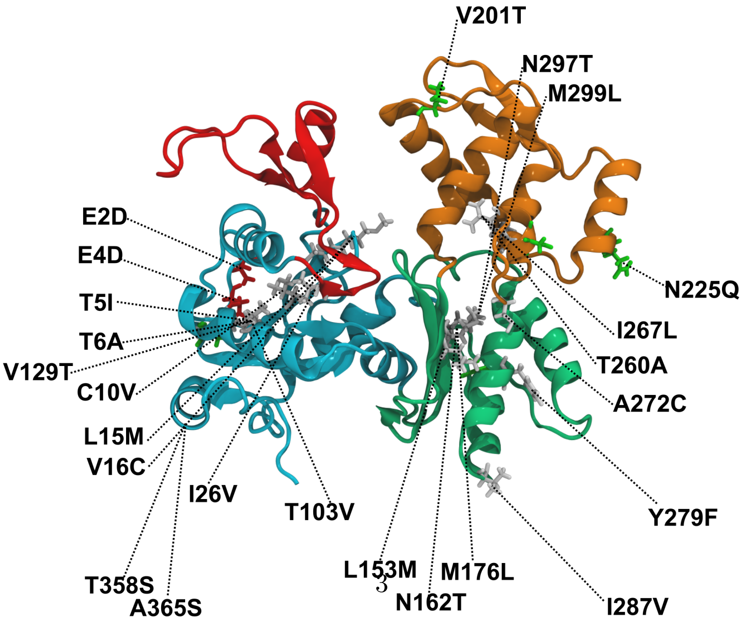


Identity:350/375 (93.3%)      Similarity:362/375 (96.5%)

**Supplementary Figure 2  Sequence alignment of $\alpha$-skeletal and $\beta$-cytoplasmic actin and positioning of mutations.** The genes for the sequence alignment correspond to ACTA1 and ACTB for the $\alpha$- and $\beta$-actin respectively. The sequence alignment has been performed using EMBOSS NEEDLES. Differences between sequences are colored based on low (blue) or high (red) chemical differences. Residues, represented in licorice, are colored according to their chemical properties (red : charged, white: apolar, green: polar). Changes are labelled with the first and last letters corresponding to the $\alpha$- and $\beta$-actin residues respectively. In addition of the histidine 73 methylation, the two first residues in the $\beta$-actin were removed and the D3 was acetylated in our model and in our experimental asssays.

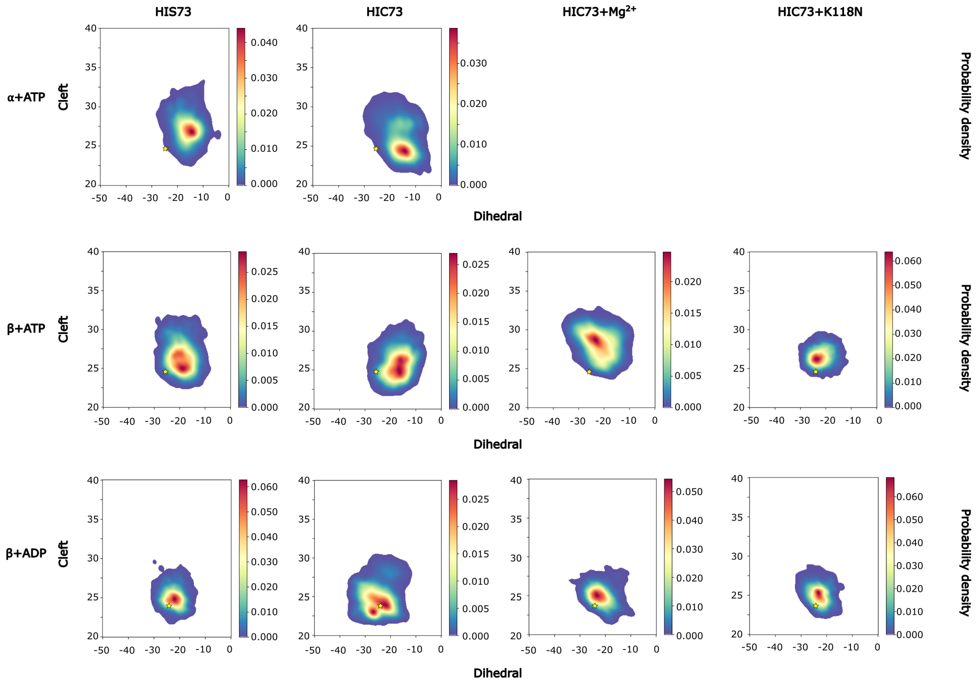


**Supplementary Figure 3 Cleft-Diedral of all monomer systems. First line:** ATP-bound $\alpha$-actin. **Second line:** ATP-bound $\beta$-actin. **Third line:** ADP-bound $\beta$-actin. Yellow star represents the starting structure, 1NWK and 1J6Z for ATP and ADP-actin respectively. Statistical significance of differences have been assessed using a two-sided Kolmogorov-Smirnov test (Table S3)

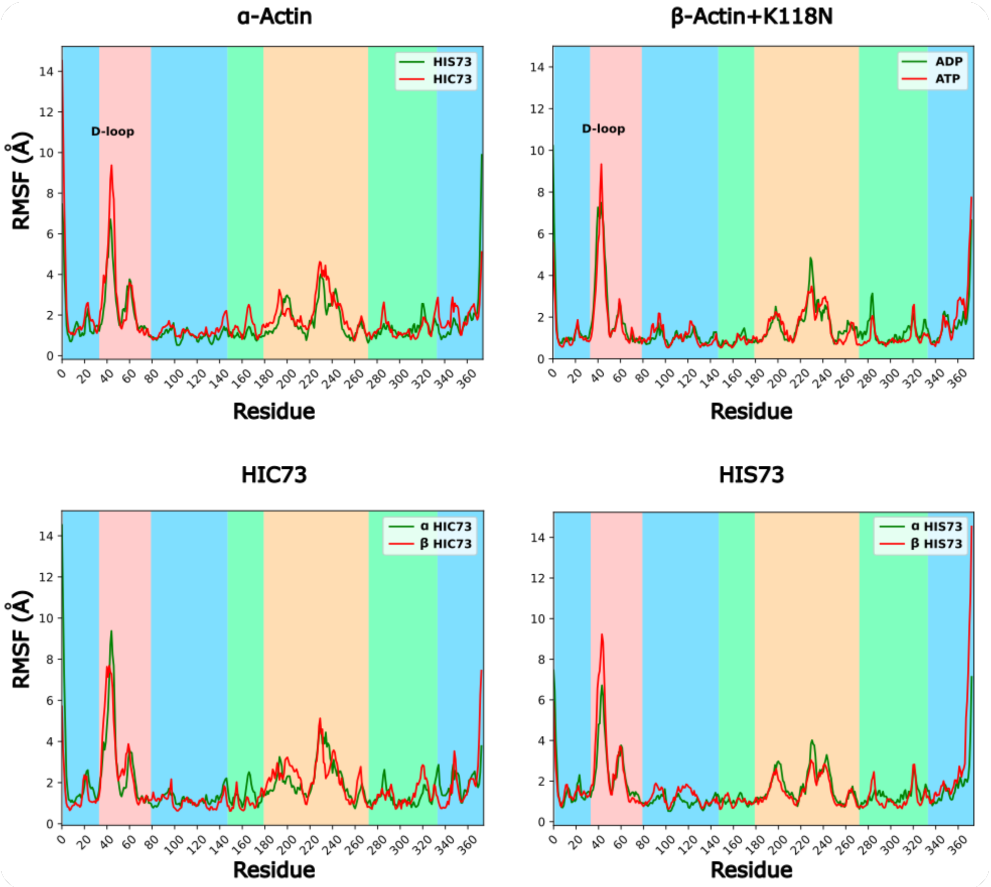


**Supplementary Figure 4 RMSF of $\alpha$-actin and $\beta$-actin + K118N systems. Left panel:** ATP-bound $\alpha$-actin non-methylated (green line) and methylated (red line). **Right panel:** ADP-bound (green line) and ATP-bound (red line) $\beta$-actin with K118N mutation.

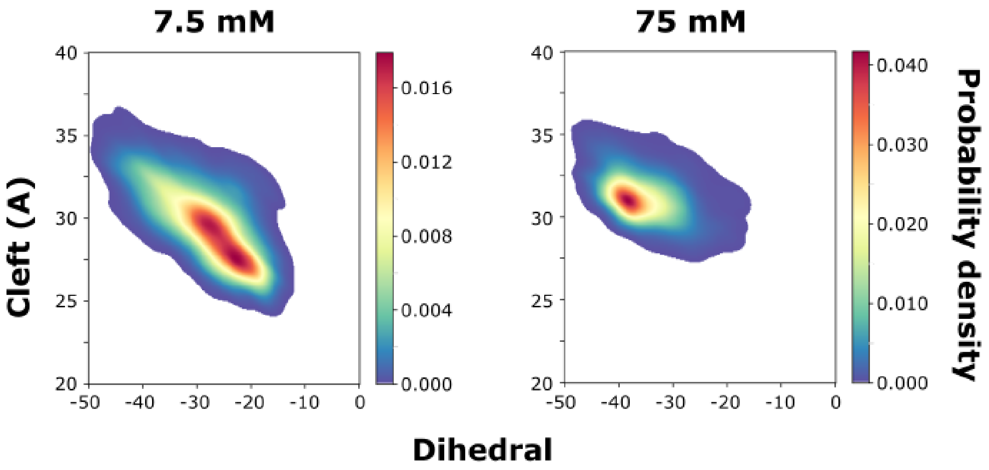


**Supplementary Figure 5 Cleft-Dihedral of 100K particle simulations at 7.5 and 75mM $MgCl_2$.** To test magnesium concentrations closer to experimental conditions, we designed systems with a larger solvatation box, reaching more than 100k particles. This allowed us to simulate a concentration of 7.5mM $MgCl_2$ (5 magnesium ions in the box). To prevent a possible size effect of the box, we also performed a simulation at 75mM $MgCl_2$ (50 magnesium ions in the box) in this larger solvation box. For these concentrations as well as for the smaller system (see Figure 1-B) we observed an opening of the cleft and an increase of the dihedral angle.

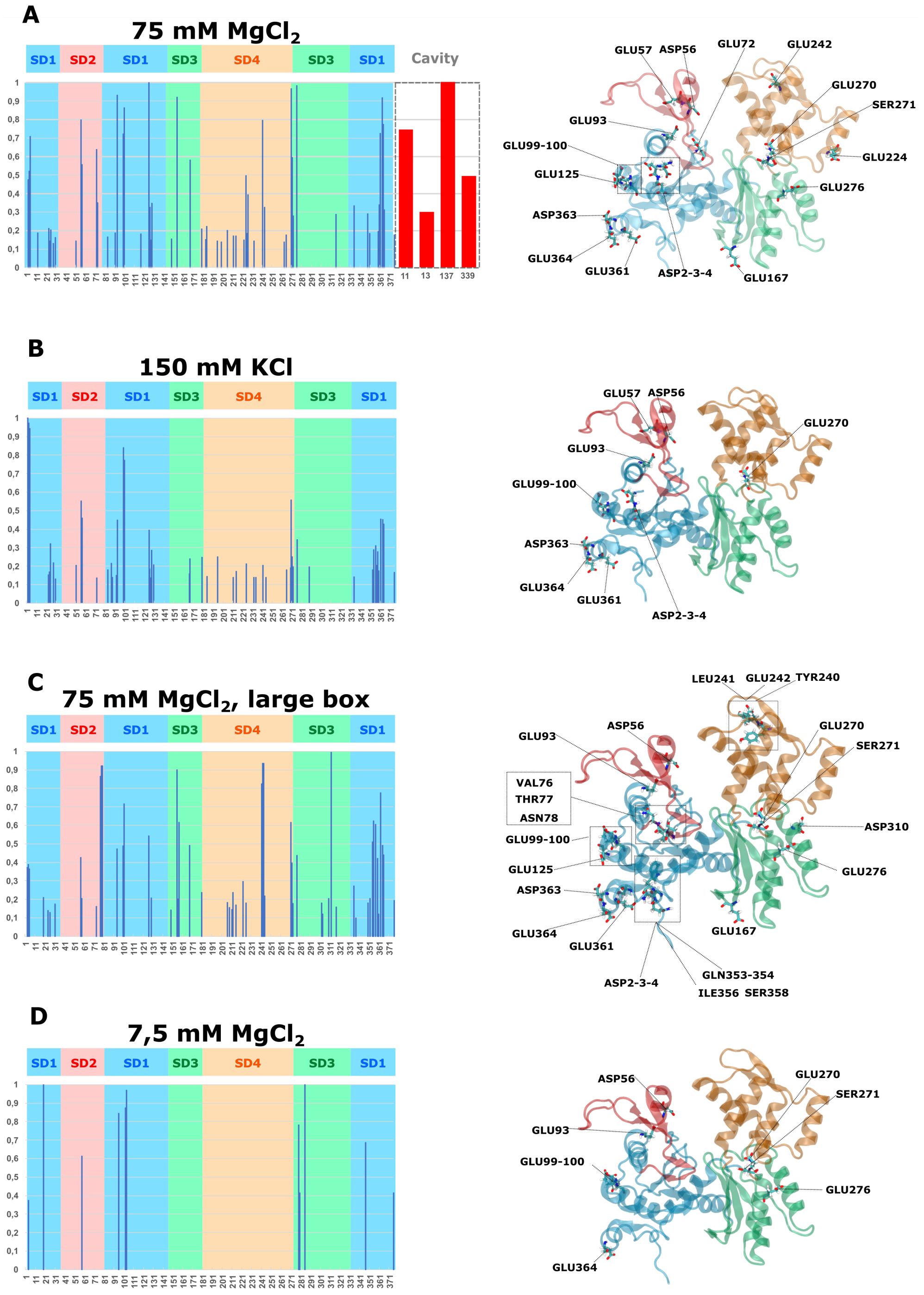


**Supplementary Figure 6 Amino acids responsible for coordinating magnesium ions. A-** Representation of normalized contacts between amino acids and Mg$^2$+ ions among all subdomains. In red normalized contacts of residues located in the cavity. **B-** For comparison, Representation of normalized contacts between amino acids and K$^+$ ions among all subdomains.**C-** Representation of normalized contacts between amino acids and Mg$^2$+ ions among all subdomains in simulation at a concentration of 75 mM Mg$^2$+, in a larger simulation box.**D-** Representation of normalized contacts between amino acids and Mg$^2$+ ions among all subdomains in simulation at a concentration of 7.5 mM Mg$^2$+, in a larger simulation box. On the right of each graph, all atom representation of amino acids with contacts frequencies above 0.4.

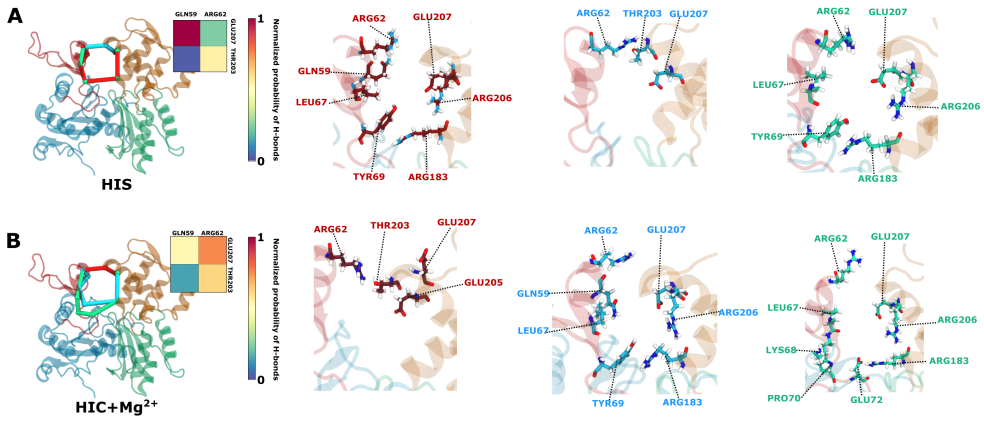


**Supplementary Figure 7 Correlations pathways in ADP-bound $\beta$-actin systems.** Representation of 3 most represented pathways of communication between ARG62 and GLU207 for **A-**HIS73, **B-**HIC73+MG systems. Concerned amino acids are represented in licorice. Insets show density of main H-bonds between SD2 and SD4 subdomains.

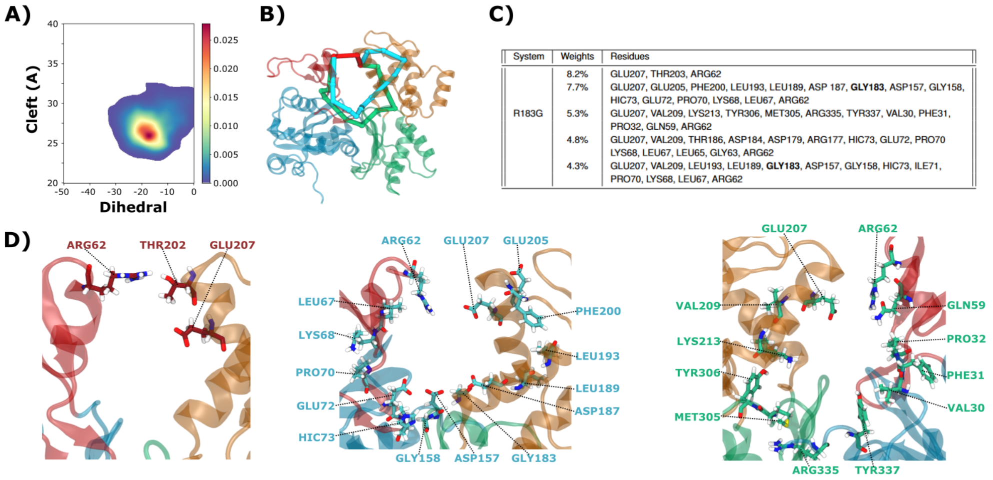


| System | Weights | Residues |
|---|---|---|
| R183G | 8.2% | GLU207, THR203, ARG62 |
| | 7.7% | GLU207, GLU205, PHE200, LEU193, LEU189, ASP 187, **GLY183**, ASP157, GLY158, HIC73, GLU72, PRO70, LYS68, LEU67, ARG62 |
| | 5.3% | GLU207, VAL209, LYS213, TYR306, MET305, ARG335, TYR337, VAL30, PHE31, PRO32, GLN59, ARG62 |
| | 4.8% | GLU207, VAL209, THR186, ASP184, ASP179, ARG177, HIC73, GLU72, PRO70 LYS68, LEU67, LEU65, GLY63, ARG62 |
| | 4.3% | GLU207, VAL209, LEU193, LEU189, **GLY183**, ASP157, GLY158, HIC73, ILE71, PRO70, LYS68, LEU67, ARG62 |

**Supplementary Figure 8 ATP-bound R183G $\beta$-actin fluctuations and pathways. A-**Cleft-Dihedral of R183G mutant in the ATP state. **B-** Representation of 3 most represented pathways of communication between ARG62 and GLU207. **C-** Reweighted percentage of 5 first shortests paths between ARG62 and GLU207. **D-** Licorice representation of first (red), second(blue) and third(green) most represented pathways.Differences observed between systems are all significant (Table S3).

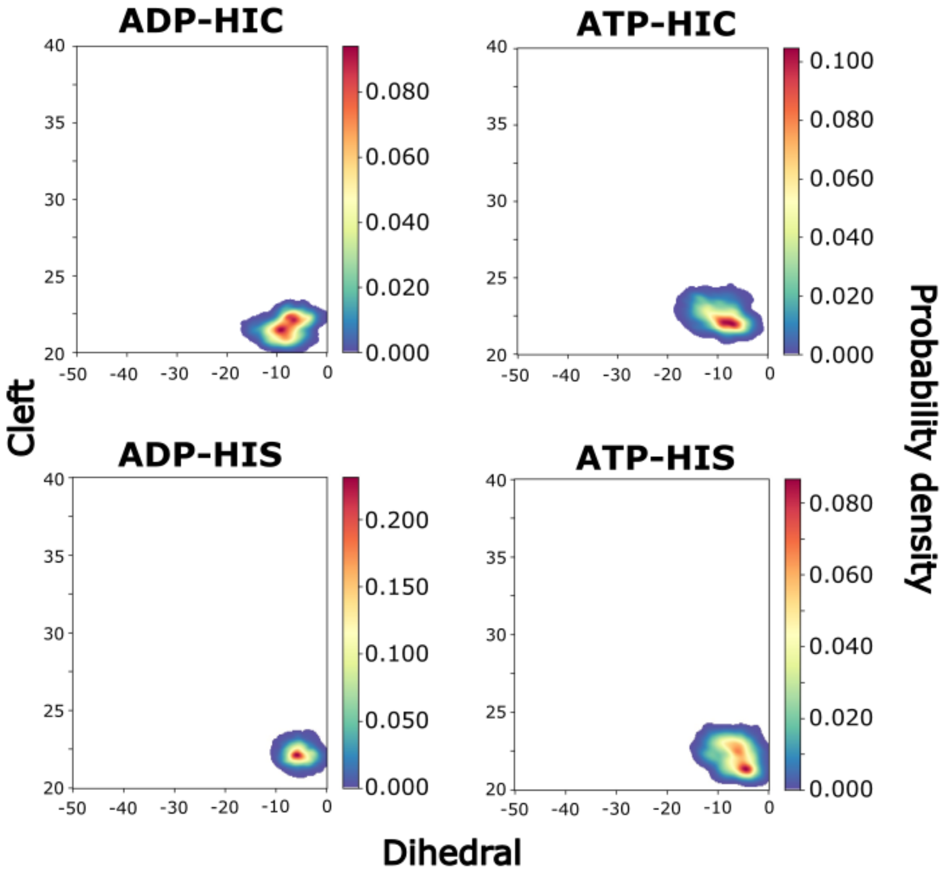


**Supplementary Figure 9** Cleft-Diedral of B-1 subunit of the 4mer systems in ADP-HIC (upper left), ATP-HIC(upper right), ADP-HIS (lower left), ATP-HIS (lower right) state. Differences observed between systems are all significant (Table S3).

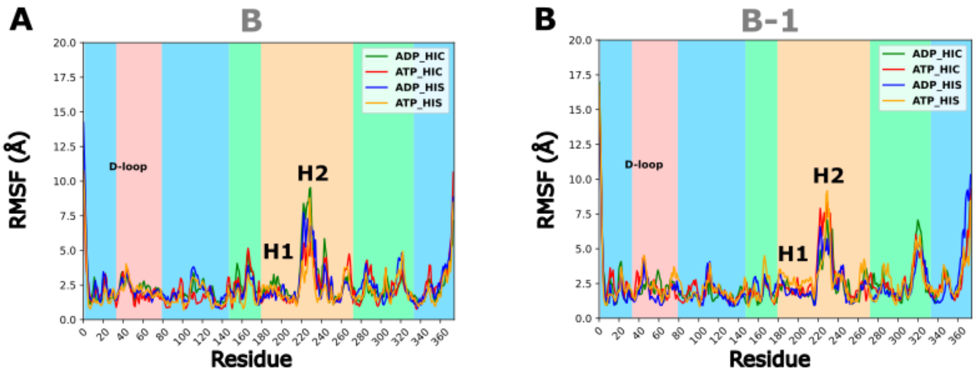


**Supplementary Figure 10 RMSF of the B and B-1 subunits of the 4mer systems in the ADP and the ATP states. A:** RMSF of the B subunits of the 4mer systems in the ADP-HIC (green line), ATP-HIC (red line), ADP-HIS (Blue line) and the ATP-HIS (Orange line) state. **B:** RMSF of the B-1 subunits of the 4mer systems in the ADP (green line) and the ATP (red line) state.

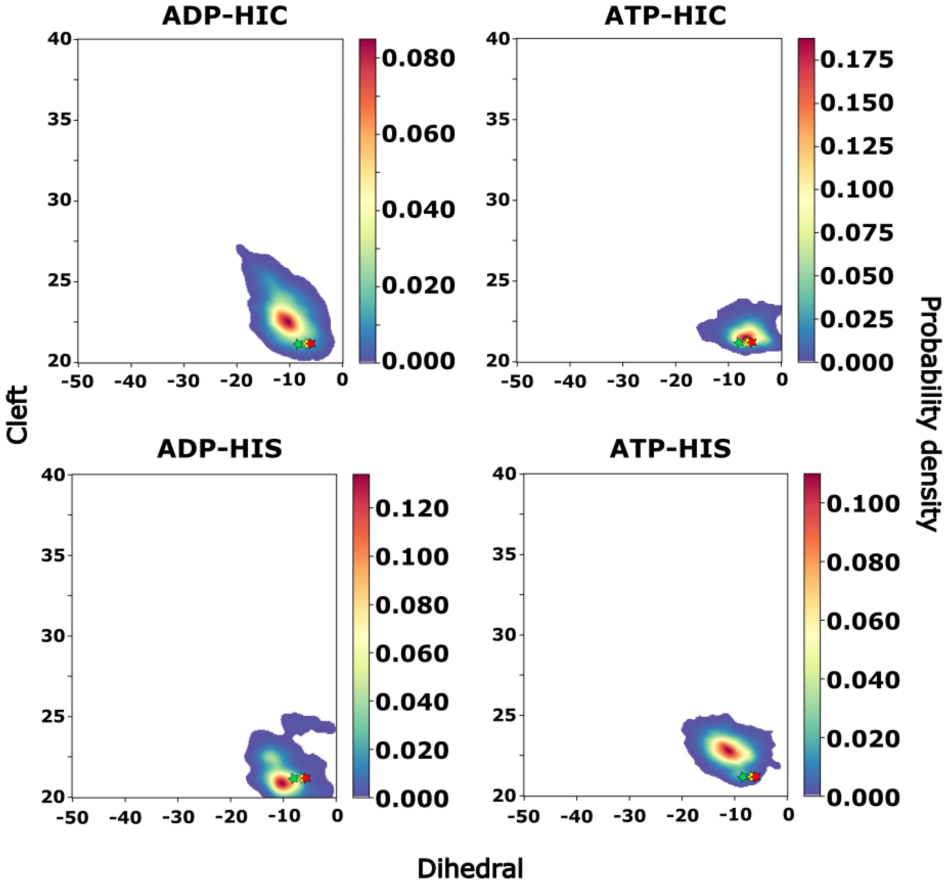


**Supplementary Figure 11 Projection of experimental structures on Cleft-Dihedral maps of the ultimate (B) subunit at the barbed end in ADP-HIC, ATP-HIC, ADP-HIS and ATP-HIS states.** The green, yellow and red stars refer respectively to the structure of Oosterheert and coauthors [1], the structure of Cartman and coauthors [2] and our starting structure [3].

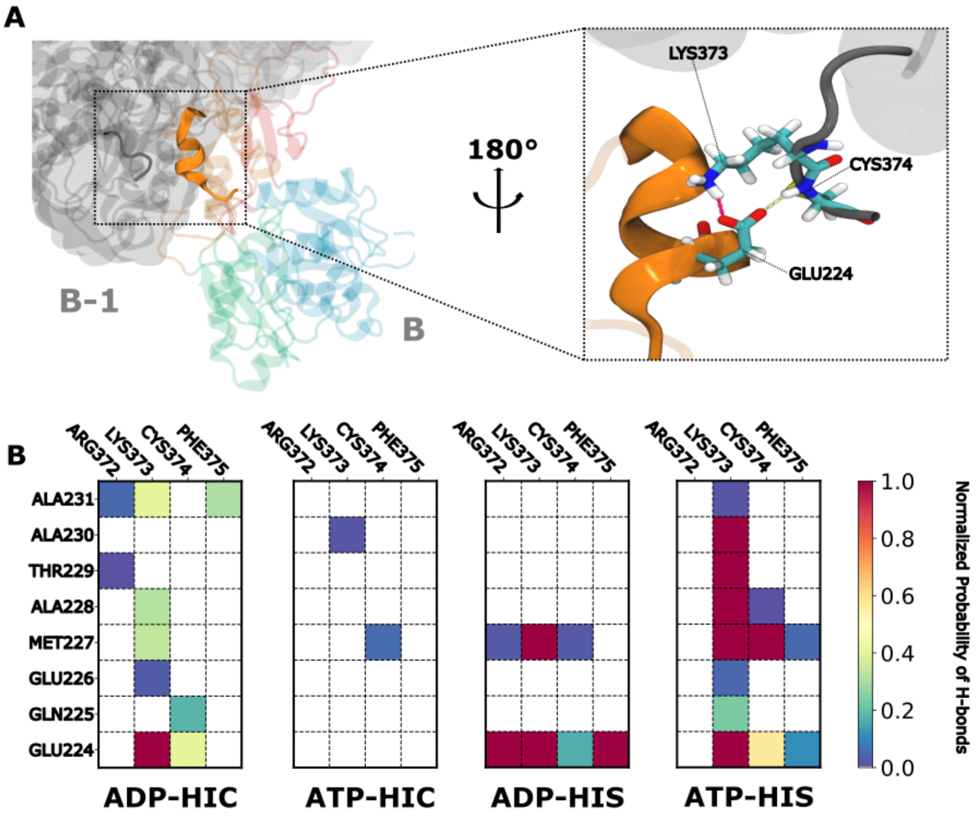


**Supplementary Figure 12 Interactions between B and B-1 monomer. A-** Highlight of C-terminal extremity of B-1 subunit and 220-230 helix of subunit B. Representation of the highest hydrogen bonds between C-terminal extremity of B-1 subunit and 220-230 helix of subunit B. **B-** Normalized probability of hydrogen bonds between C-terminal extremity of B-1 subunit and 220-230 helix of subunit B in ADP-HIC, ATP-HIC, ADP-HIS and ATP-HIC states

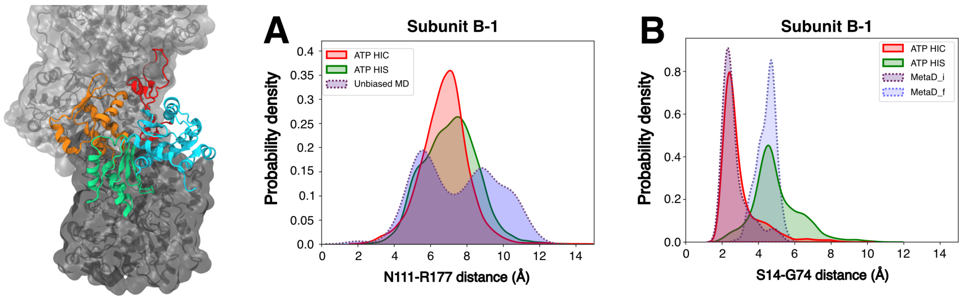


**Supplementary Figure 13** **A-** Distribution of distances between residues N111 and R177 in ATP-HIS and ATP-HIC for the penultimate (B-1) actin monomer at the barbed end. The unbiased MD curve corresponds to distances extracted from unbiaised MD simulations from [1]. **B-** Distribution of distances between residues S14 and G74 in ATP-HIS and ATP-HIC for the penultimate (B-1) actin monomer at the barbed end. The metaD_i (purple) and metaD_f (blue) distributions correspond respectively to distances observed in the initial and final states of metadynamics simulations extracted from ref [1] (see Method). In this simulation the G74-S14 backdoor is closed in the first half of the simulation(metaD_i, purple dashed curve), and open in the second half of the simulation (metaD_f blue dashed curve).

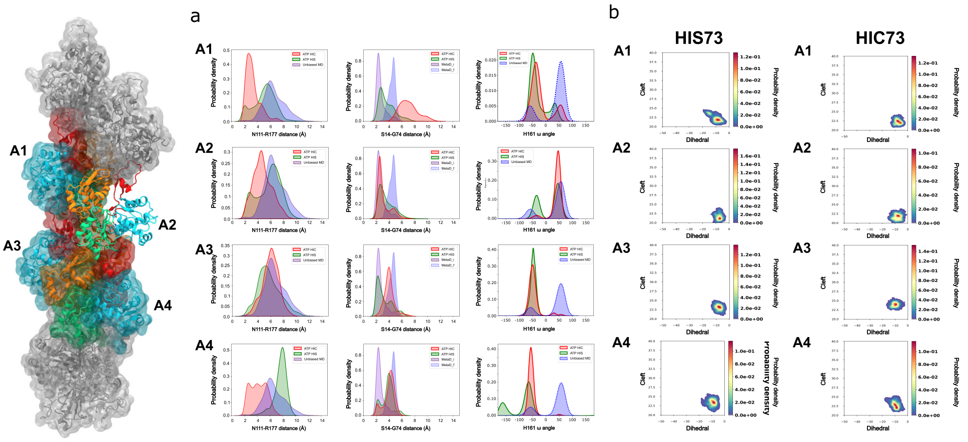


**Supplementary Figure 14 Analysis of cleft-dihedral and backdoors opening in the filament.** **A-** For the N111-R1777 backdoor, the blue distribution correspond to distances observed in unbiased simulations extracted from [1] for subunit B. For the S14-G74 backdoor, the metaD_i (purple) and metaD_f (blue) distributions correspond respectively to distances observed in the initial and final states of metadynamics simulations extracted from ref [1] (see Method). In this simulation the G74-S14 backdoor is closed in the first half of the simulation(metaD_i, purple dashed curve), and open in the second half of the simulation (metaD_f blue dashed curve). **B-** Cleft-Dihedral maps for the different actin subunits for unmethylated (HIS) and methylated (HIC) histdine 73.

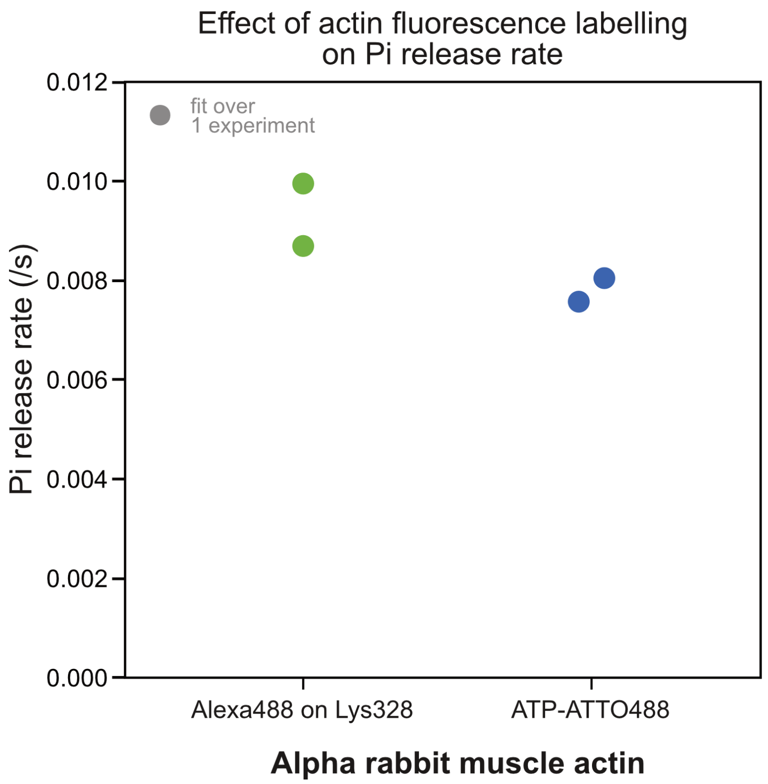


**Supplementary Figure 15 Effect of fluorescence labelling on Pi release rate for $\alpha$-skeletal actin filaments.** Pi-release rate measured from $\alpha$-skeletal actin filament depolymerization assays, as represent in figure 5F. Each measurement data point is the value measured from the acceleration of the depolymerization rate of at least 30 individual filaments. Actin was either 10%-labelled on surface Lysine 328 with Alexa488 fluorophore, or labelled using 0.5 µM ATP-Atto488 in the polymerization buffer (see Methods).

**Supplementary Movie 1** Morphing between the two structures of unmethylated $\beta$-actin corresponding to green and blue points in figure S1

**Supplementary Movie 2** Morphing between the two structures of methylated $\beta$-actin corresponding to green and blue points in figure S1

**Supplementary Movie 3** Morphing between the two structures of methylated $\beta$-actin with a 75mM $Mg^{2+}$ concentration corresponding to green and blue points in figure S1

**Supplementary Table 1 systems modelled:**. The different columns correspond to actin isoform, post-translational modification (P.T.M.) of histidine 73 (HIS: unmodified and HIC: methylated), nucleotide type, ions concentrations, type of system (Mono: monomer; number of atoms in parenthesis), and initial actin structure (G: globular actin, F: filament).

| **Seq.** | **P.T.M.** | **Nuc.** | **Ions (mM)** | **System** (atoms) | **Struc.** |
|---|---|---|---|---|---|
| $\alpha$ | HIS | ATP | 150 $KCl$ | Mono (~55k) | G |
| $\alpha$ | HIC | ATP | 150 $KCl$ | Mono (~55k) | G |
| $\beta$ | HIS | ATP | 150 $KCl$ | Mono (~55k) | G |
| $\beta$ | HIC | ATP | 150 $KCl$ | Mono (~55k) | G |
| $\beta$ | HIC | ATP | 75 $MgCl_2$ | Mono (~55k) | G |
| $\beta$ | HIC | ATP | 75 $MgCl_2$ | Mono (~100k) | G |
| $\beta$ | HIC | ATP | 7.5 $MgCl_2$ | Mono (~100k) | G |
| $\beta$ | HIS | ADP | 150 $KCl$ | Mono (~55k) | G |
| $\beta$ | HIC | ADP | 150 $KCl$ | Mono (~55k) | G |
| $\beta$ | HIC | ADP | 75 $MgCl_2$ | Mono (~55k) | G |
| $\beta_{R183G}$ | HIC | ATP | 150 $KCl$ | Mono (~55k) | G |
| $\beta_{K118N}$ | HIC | ATP | 150 $KCl$ | Mono (~55k) | G |
| $\beta_{K118N}$ | HIC | ADP | 150 $KCl$ | Mono (~55k) | G |
| $\beta$ | HIC | ATP | 150 $KCl$ | 4-mer (~250k) | F |
| $\beta$ | HIC | ADP | 150 $KCl$ | 4-mer (~250k) | F |
| $\beta$ | HIS | ATP | 150 $KCl$ | 4-mer (~250k) | F |
| $\beta$ | HIS | ADP | 150 $KCl$ | 4-mer (~250k) | F |
| $\beta$ | HIC | ATP | 150 $KCl$ | 8-mer (~525k) | F |
| $\beta$ | HIS | ATP | 150 $KCl$ | 8-mer (~525k) | F |

| System | Weights | Residues |
|---|---|---|
| ATP+HIS | 19.0% | GLU207, ARG62 |
| | 7.9% | GLU207, THR203, ARG62 |
| | 7.1% | GLU207, ARG206, ARG183, GLU72, PRO70, LYS68, LEU67, GLN59, ARG62 |
| | 5.5% | GLU207, ARG206, ARG183, ASP157, GLY156, HIS73, GLU72, TYR69, LEU67, ARG62 |
| | 4.2% | GLU207, ARG206, ARG183, TYR69, LYS68, LEU67, GLN59, ARG62 |
| ATP+HIC | 8.9% | GLU207, ARG206, ARG183, GLU72, TYR69, LEU67, GLN59, ARG62 |
| | 8.3% | GLU207, ARG206, ARG183, GLU72, PRO70, LYS68, LEU67, GLN59, ARG62 |
| | 5.5% | GLU207, ARG206, ARG183, TYR69, LEU67, GLN59, ARG62 |
| | 5.1% | GLU207, VAL209, LEU189, ASP187, ARG183, TYR69, LEU67, GLN59, ARG62 |
| | 4.9% | GLU207, VAL209, LYS213, GLY182, ARG183, GLU72, PRO70, LYS68, THR66, LEU65, LYS61, ARG62 |
| ATP+HIC+$Mg^{2+}$ | 9.3% | GLU207, VAL209, LYS213, THR203, MET305, LYS336 , TYR337, PRO27, VAL30, PHE31, PRO32, GLY55, GL59, ARG62 |
| | 5.7% | GLU207, ARG206, ARG183, ASP157, GLY158, HIC73, ILE71, GLU72, TYR69, LEU67, GLN59, ARG62 |
| | 4.6% | GLU207, ARG62 |
| | 4.1% | GLU207, VAL209, LEU189, ASP187, ARG183, GLU72, TYR69, LYS68, LEU67, GLN59, ARG62 |
| | 4.1% | GLU207, ARG206, LEU188, ASP187, ARG183, GLU72, TYR69, LYS68, LEU67, GLN59, ARG62 |
| ADP+HIS | 18.8% | GLU207, ARG206, ARG183, TYR69, LEU67, GLN59, ARG62 |
| | 11.2% | GLU207, ARG206, ARG183, TYR69, LEU67, ARG62 |
| | 7.5% | GLU207, THR203, ARG62 |
| | 6.3% | GLU207, ARG206, ARG183, GLU72, TYR69, LEU67, ARG62 |
| | 6.1% | GLU207, ARG206, THR186, ARG183, TYR69, LYS68, LEU67, GLN59, ARG62 |
| ADP+HIC | 25.7% | GLU207, THR203, ARG62 |
| | 10.8% | GLU207, ARG206, ARG183, TYR69, LEU67, GLN59, ARG62 |
| | 10.5% | GLU207, ARG206, ARG183, TYR69, LEU67, ARG62 |
| | 7.2% | GLU207, ARG206, ARG183, ASP157, GLY15, PRO32, ILE33, GLN59, ARG62 |
| | 6.2% | GLU207, ARG206, ARG183, TYR69, LYS68, LEU67, GLN59, ARG62 |
| ADP+HIC+$Mg^{2+}$ | 14.1% | GLU207,GLU205, THR203, ARG62 |
| | 11.3% | GLU207, ARG206, ARG183, TYR69, LEU67, GLN59, ARG62 |
| | 10.0% | GLU207, ARG206, ARG183, GLU72, PRO70, LYS68, LEU67, ARG62 |
| | 7.4% | GLU207, THR203, ARG62 |
| | 7.4% | GLU207, ARG206, ARG183, GLU72, TYR69, LEU67, GLN59, ARG62 |

**Supplementary Table 2** Reweighted percentage of 5 first shortests paths between ARG62 and GLU207.

**Supplementary Table 3** Differences observed between the measures calculated in the different systems, using a Kolmogorov-Smirnov test. To apply this test, we considered each seed of 10 ns as an independent simulation and took the data from the last frame of each seed leading to a total of 152 independent measures per simulation. * = p-value $< 0.05$. *** = p-value $< 0.001$. ns = $p > 0.05$ .

| | | | | |
|---|---|---|---|---|
| **FIG1** | **Cleft values** | **HIS** | **HIC** | * |
| | | **HIS** | **HIC_MG** | *** |
| | | **HIC** | **HIC_MG** | *** |
| | **Diedral values** | **HIS** | **HIC** | *** |
| | | **HIS** | **HIC_MG** | *** |
| | | **HIC** | **HIC_MG** | *** |
| **FIG2** | **Cavity volume** | **HIS** | **HIC** | *** |
| | | **HIS** | **HIC_MG** | *** |
| | | **HIC** | **HIC_MG** | ns |
| | **Distance Mg-Pg** | **HIS** | **HIC** | * |
| | | **HIS** | **HIC_MG** | *** |
| | | **HIC** | **HIC_MG** | * |
| | **Number of water molecules** | **HIS** | **HIC** | ns |
| | | **HIS** | **HIC_MG** | ns |
| | | **HIC** | **HIC_MG** | ns |
| **FIG4** | **Cleft values** | **HIS** | **HIC** | *** |
| | | **HIS** | **HIC_MG** | *** |
| | | **HIC** | **HIC_MG** | *** |
| | **Diedral values** | **HIS** | **HIC** | *** |
| | | **HIS** | **HIC_MG** | *** |
| | | **HIC** | **HIC_MG** | *** |
| | **Cavity volume** | **HIS** | **HIC** | ** |
| | | **HIS** | **HIC_MG** | *** |
| | | **HIC** | **HIC_MG** | *** |
| | **Distance Mg-Pb** | **HIS** | **HIC** | ns |
| | | **HIS** | **HIC_MG** | *** |
| | | **HIC** | **HIC_MG** | *** |
| | **Number of water molecules** | **HIS** | **HIC** | *** |
| | | **HIS** | **HIC_MG** | *** |
| | | **HIC** | **HIC_MG** | *** |
| **FIG5** | **Cleft values** | **ATP-HIS** | **ATP-HIC** | *** |
| | | **ATP-HIS** | **ADP-HIS** | *** |
| | | **ATP-HIS** | **ADP-HIC** | *** |
| | | **ATP-HIC** | **ADP-HIS** | *** |
| | | **ATP-HIC** | **ADP-HIC** | *** |
| | | **ADP-HIS** | **ADP-HIC** | *** |
| | **Diedral values** | **ATP-HIS** | **ATP-HIC** | *** |
| | | **ATP-HIS** | **ADP-HIS** | ns |
| | | **ATP-HIS** | **ADP-HIC** | *** |
| | | **ATP-HIC** | **ADP-HIS** | *** |
| | | **ATP-HIC** | **ADP-HIC** | *** |
| | | **ADP-HIS** | **ADP-HIC** | *** |
| **FIG SUPP 3** | **Cleft values** | $\alpha$**-HIS** | $\alpha$**-HIC** | *** |
| | | $\alpha$**-HIS** | $\beta$**-HIS** | *** |
| | | $\alpha$**-HIC** | $\beta$**-HIC** | *** |
| | | **ATP-K118N** | **ATP-**$\beta$ | *** |
| | | **ADP-K118N** | **ADP-**$\beta$ | *** |
| | | **ATP-K118N** | **ADP-K118N** | *** |
| | **Diedral values** | $\alpha$**-HIS** | $\alpha$**-HIC** | *** |
| | | $\alpha$**-HIS** | $\beta$**-HIS** | *** |
| | | $\alpha$**-HIC** | $\beta$**-HIC** | *** |
| | | **ATP-K118N** | **ATP-**$\beta$ | *** |
| | | **ADP-K118N** | **ADP-**$\beta$ | *** |
| | | **ATP-K118N** | **ADP-K118N** | *** |

### References

[1] Oosterheert, W., Blanc, F.E.C., Roy, A., Belyy, A., Sanders, M.B., Hofnagel, O., Hummer, G., Bieling, P., Raunser, S.: Molecular mechanisms of inorganic-phosphate release from the core and barbed end of actin filaments. Nature Structural & Molecular Biology (2023) https://doi.org/10.1038/s41594-023-01101-9

[2] Carman, P.J., Barrie, K.R., Rebowski, G., Dominguez, R.: Structures of the free and capped ends of the actin filament. Science **380**(6651), 1287–1292 (2023) https://doi.org/10.1126/science.adg6812 https://www.science.org/doi/pdf/10.1126/science.adg6812

[3] Gurel, P.S., Kim, L.Y., Ruijgrok, P.V., Omabegho, T., Bryant, Z., Alushin, G.M.: Cryo-EM structures reveal specialization at the myosin VI-actin interface and a mechanism of force sensitivity. eLife **6**(Md), 1–33 (2017) https://doi.org/10.7554/eLife.31125
